# Leveraging non-structural data to predict structures of protein–ligand complexes

**DOI:** 10.1101/2020.06.01.128181

**Authors:** Joseph M. Paggi, Julia A. Belk, Scott A. Hollingsworth, Nicolas Villanueva, Alexander S. Powers, Mary J. Clark, Augustine G. Chemparathy, Jonathan E. Tynan, Thomas K. Lau, Roger K. Sunahara, Ron O. Dror

**Affiliations:** Department of Computer Science, Stanford University, Stanford, CA 94305, USA; Department of Molecular and Cellular Physiology, Stanford University School of Medicine, Stanford, CA 94305, USA; Department of Structural Biology, Stanford University School of Medicine, Stanford, CA 94305, USA; Institute for Computational and Mathematical Engineering, Stanford University, Stanford, CA 94305, USA; Department of Chemistry, Stanford University, Stanford, CA 94305, USA; Department of Pharmacology, University of California San Diego School of Medicine, La Jolla, CA 92093, USA

## Abstract

Over the past fifty years, tremendous effort has been devoted to computational methods for predicting properties of ligands that bind macromolecular targets, a problem critical to rational drug design. Such methods generally fall into two categories: physics-based methods, which directly model ligand interactions with the target given the target’s three-dimensional (3D) structure, and ligand-based methods, which predict ligand properties given experimental measurements for similar ligands. Here we present a rigorous statistical framework to combine these two sources of information. We develop a method to predict a ligand’s pose—the 3D structure of the ligand bound to its protein target—that leverages a widely available source of information: a list of other ligands that are known to bind the same target but for which no 3D structure is available. This combination of physics-based and ligand-based modeling improves upon state-of-the-art pose prediction accuracy across all major families of drug targets. As an illustrative application, we predict binding poses of antipsychotics and validate the results experimentally. Our statistical framework and results suggest broad opportunities to predict diverse ligand properties using machine learning methods that draw on physical modeling and ligand data simultaneously.

## Introduction

Binding of small-molecule ligands to proteins is one of the most fundamental processes in biology, and the great majority of drugs are ligands that exert their effects by binding to a target protein. Predicting properties of protein–ligand interactions—including three-dimensional (3D) structures, binding affinities, binding kinetics, selectivity, and functional effects—is critical both to the rational design of effective medicines and to solving major problems in molecular biology. An enormous amount of work has thus focused on the development of computational methods to predict these properties (1, 2).

Such computational methods generally fall into two categories. “Physics-based” approaches use a 3D structure of the target protein and exploit an understanding of the physics of protein–ligand interactions (3). “Ligand-based” approaches use experimental measurements for many ligands of a given property (e.g., affinity) at a given target and employ pattern matching to predict the corresponding property for other ligands (4, 5).

Can one combine these two paradigms, and the orthogonal sources of information they leverage, in a systematic, principled manner? This has proven challenging, particularly when making predictions for ligands substantially different from those for which experimental data is available. It is especially difficult when one wishes to predict properties different from those measured experimentally—e.g., to predict ligand properties that are difficult to determine experimentally by exploiting experimental data that is easy to collect.

Here we present a method, ComBind, that overcomes these obstacles to substantially improve prediction of a ligand’s binding pose at a given target protein. Determining a ligand’s binding pose—the three-dimensional coordinates of the ligand’s atoms when bound to the target—is critical to structure-based optimization of the ligand’s pharmacological properties, as well as to understanding how it influences its target. Indeed, knowledge of a ligand’s binding pose is so advantageous that researchers in industry and academia often spend months or years to solve an experimental structure of a particular ligand in complex with a target protein.

Because experimental structure determination is time-consuming, expensive, and sometimes impossible, tremendous effort has been invested in the development of *in silico* “docking” methods for predicting ligand binding poses (6–15). These are physics-based approaches: given a structure of the target protein, they sample many candidate poses of a ligand and rank these poses using scoring functions that approximate the energetic favorability of each pose, typically by capturing interatomic interactions such as hydrogen bonds and van der Waals forces (**Fig. 1A**). Despite the development of dozens of docking software packages over the past 40 years, binding pose predictions are typically correct less than half the time for ligands substantially different from those in the experimental structures used for docking (**Supplementary Table 1**).

**Figure 1:**
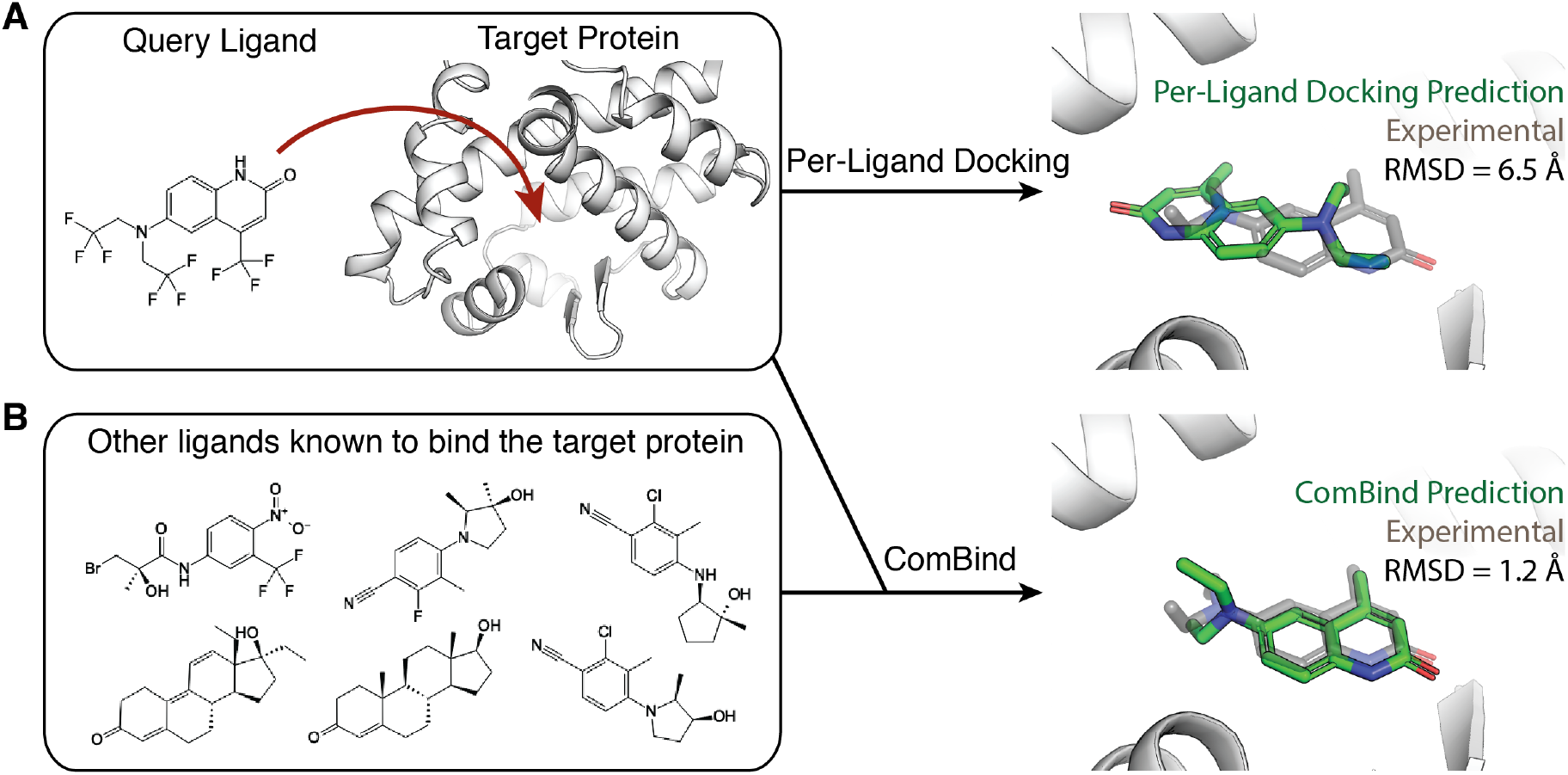
ComBind leverages non-structural data to improve ligand binding pose predictions. (A) Standard docking methods take as input the chemical structure of the query ligand and the 3D structure of the target protein and predict a binding pose using a per-ligand scoring function. (B) ComBind additionally considers other ligands known to bind the target protein (whose binding poses are not known), resulting in more accurate predictions. For clarity, hydrogen and fluorine atoms are not shown in the 3D renderings.

ComBind improves binding pose prediction by exploiting a widely available type of non-structural data: the identities of other ligands known to bind the same target (**Fig. 1B**). Collecting such data is typically far easier than structure determination. Indeed, such data is routinely collected in drug development campaigns and is already available in public databases such as ChEMBL for the vast majority of recognized drug targets (16).

How can a list of other ligands that bind to the target protein—but whose binding poses are unknown—be used to improve pose prediction? Medicinal chemists have long recognized that distinct ligands tend to bind a given protein in similar poses; even ligands sharing no common substructure often form similar interactions with the target protein (**Fig. 2A**). This intuition has a sound basis in physics. For example, the energetic favorability of a protein–ligand hydrogen bond depends on the mobility of the protein atoms involved and their ability to form hydrogen bonds with water in the absence of the ligand (17). These factors contribute similarly to binding of different ligands but are difficult to predict from a static protein structure alone.

**Figure 2:**
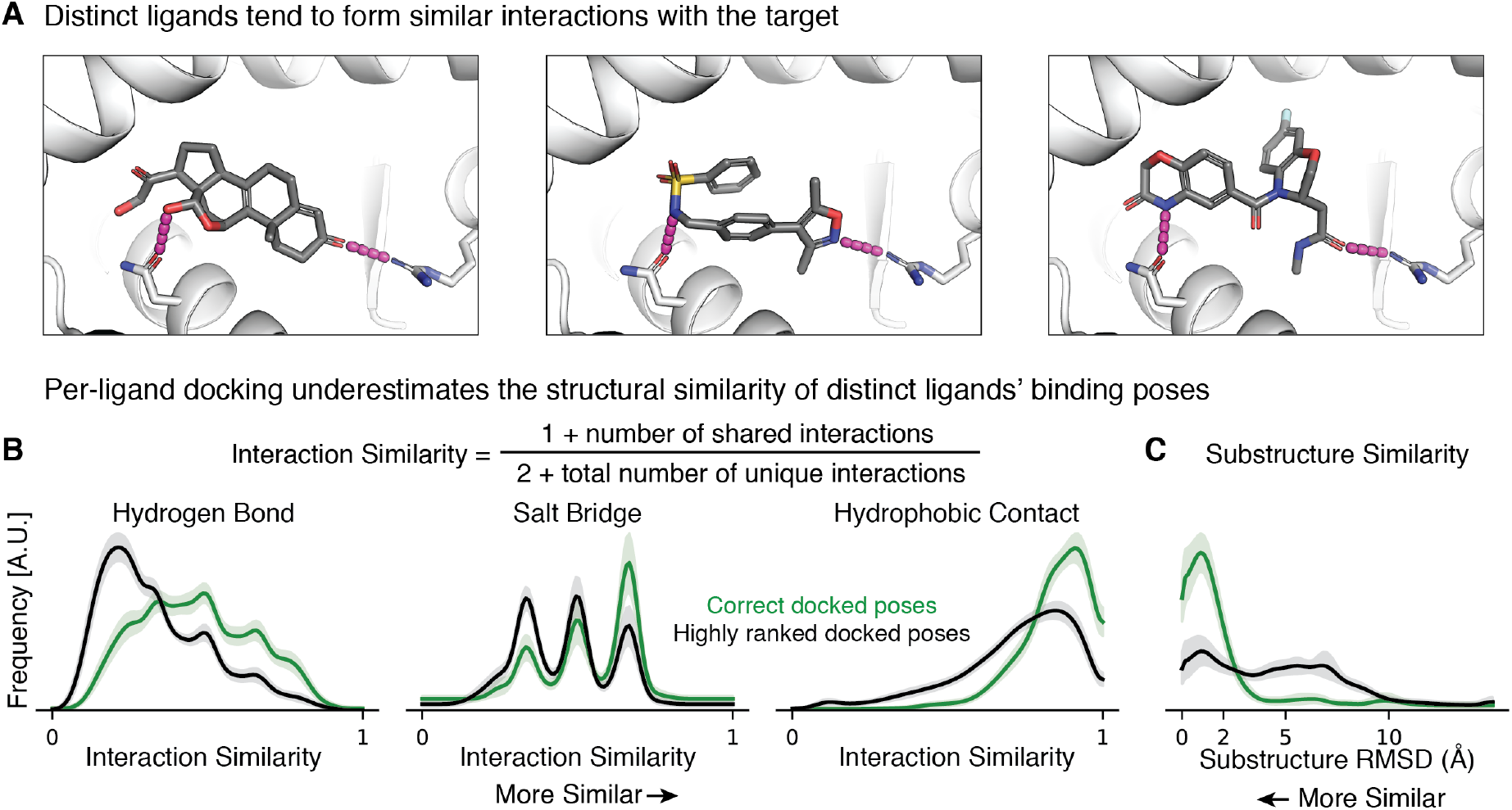
Distinct ligands that bind to a given target protein often adopt similar binding poses and do so more frequently than predicted by a state-of-the-art per-ligand docking method. (A) Chemically distinct ligands share key interactions with the mineralocorticoid receptor (PDB IDs: 2AA2, 5L7E, 5MWP). (B) Across a set of 3115 ligand pairs, interaction similarities are generally higher in pairs of correct poses than in pairs including all poses ranked highly by a per-ligand scoring function. Shading depicts the per-target standard error of the mean. (C) Across a set of 690 ligand pairs with a shared substructure, the substructure tends to be placed more similarly in correct poses than in other poses ranked highly by a per-ligand scoring function. Substructure similarity for a pose pair was defined as the root mean square deviation (RMSD) between the largest substructure shared by the ligands (see Methods).

To develop ComBind, we first use a large set of experimentally determined structures to quantify the medicinal chemist’s intuition—in particular, to determine the probability that binding poses for different ligands will share various features. We use the results to define a scoring function that predicts the favorability of a set of binding poses comprising one pose for each ligand known to bind the target protein. By contrast to the ComBind scoring function, scoring functions typically utilized by docking software assign a score to the pose of a single ligand at a time; we thus refer to them as per-ligand scoring functions. The ComBind scoring function takes into account the similarities and differences between the poses of different ligands as well as the energetic favorability of each ligand’s pose, as evaluated by a per-ligand scoring function.

We benchmark ComBind by comparing its results to 245 experimentally determined ligand binding poses across 30 proteins representing all major families of drug targets. ComBind improves the pose prediction accuracy of state-of-the-art docking software for all major drug target families. For G-protein-coupled receptors (GPCRs), which are of particular interest both because they represent the largest family of drug targets and because their structures are notoriously difficult to determine experimentally, ComBind selects a correct binding pose over 60% more frequently, increasing the probability of correct prediction from 47% to 76%. ComBind’s results could be improved further by utilizing proprietary data generated as part of a drug discovery project (e.g., additional ligands found to bind the target).

We also illustrate the use of ComBind to predict the previously unknown binding poses of several antipsychotics at the D_2_ dopamine receptor (D_2_R), an important drug target for which experimental structure determination has proven difficult. We validate ComBind’s predictions— which differ from those of state-of-the-art docking software—using mutagenesis experiments. The results reveal structural motifs determining the potency and subtype-selectivity of D_2_R-targeted drugs.

ComBind provides a rigorous statistical foundation for combining physics-based structural modeling of protein-ligand interactions with inference based on experimental data for other ligands. It effectively leverages data on ligands that share no common scaffold or substructure with the ligand whose pose is predicted. It allows prediction of a difficult-to-determine ligand property based on a completely different type of data for other ligands. This method thus suggests a broad range of possibilities for combining physics-based and ligand-based approaches to improve prediction of various ligand properties by exploiting diverse sources of data.

## Results

### Overview of method

Given several ligands known to bind a target, ComBind solves for all their binding poses simultaneously. We use per-ligand docking software to generate a list of candidate poses for each ligand. ComBind then selects a set of poses—one for each ligand—that optimizes the ComBind scoring function. The ComBind scoring function considers both the favorability of each ligand’s pose, as assessed by a per-ligand scoring function, and the likelihood that a set of poses sharing a given level of similarity will be correct or incorrect.

Here we use the commercial docking software package Glide (9, 10) to generate candidate poses and assign a per-ligand score to each. We selected Glide because it is widely used in the pharmaceutical industry and because it ranks among the most accurate docking packages in comparative studies (18, 19). We emphasize, however, that the ComBind approach can utilize any per-ligand scoring function and pose sampling strategy, including those implemented in any per-ligand docking package.

### Quantifying the similarity of binding poses for distinct ligands

We begin by quantifying the medicinal chemist’s intuition that different ligands tend to adopt structurally similar poses when binding the same target protein. We wish not only to measure the similarity of correct poses of different ligands, but also to compare the similarity of correct poses to that of other poses ranked highly by per-ligand docking software. We consider two notions of similarity: similarity of protein–ligand interactions and similarity in position of common ligand substructures.

We compiled a set of 385 protein–ligand complex structures for 28 target proteins representing all major classes of small-molecule drug targets (**Supplementary Table 2**) (20). We docked each of the ligands using Glide and selected the 100 most highly ranked poses for each ligand. To reflect practical application of docking, we docked each ligand into an experimental structure solved in the presence of a ligand distinct from any of those being docked (“cross-docking”; see Methods). Our docking procedure did not utilize the experimentally determined poses of ligands in any way.

For all pairs of ligands for each target protein, we compute the similarity between each pose of one ligand and each pose of the other ligand. We use this data to calculate a probability distribution over similarity values; we refer to this distribution as the reference distribution.

We also compute similarities between each pair of correct poses (again, one pose per ligand), where a pose is considered correct if it is within 2.0 Å root mean squared deviation (RMSD) to the experimentally determined pose. We use these data to calculate a second probability distribution over similarity values, the native distribution. When calculating the native distribution, we use correct poses from the lists generate by Glide instead of using the experimentally determined poses directly, such that the similarity statistics we calculate will be most applicable to candidate poses considered during computational pose prediction.

We evaluate pose similarity separately for different types of protein–ligand interactions: hydrogen bonds, salt bridges, and hydrophobic contacts (**Fig. 2B, Supplementary Fig. 1A**). Given a pair of poses, we evaluate the similarity for each interaction type by cataloging the set of protein residues with which each ligand forms an interaction of the given type and then comparing the sizes of the intersection and union of these sets. Their ratio (the Tanimoto coefficient (21)) increases when shared interactions are formed and decreases when either ligand forms an unshared interaction. To make this metric well-defined when neither ligand forms any interactions of a particular type, we add pseudo counts. For all interaction types, the native distribution exhibits higher similarity than the reference distribution—that is, pairs of correct poses form more similar interactions than other pairs of poses ranked highly by the per-ligand scoring function (**Fig. 2B**).

We define substructure similarity as the RMSD of atom positions of the largest chemical substructure shared by a pair of ligands (**Supplementary Fig. 1B**). We evaluated substructure similarity for pairs of ligands that shared a substructure at least half the size of the smaller ligand. For this similarity metric, too, the native distribution exhibits higher similarity than the reference distribution, indicating that the common substructure tends to be more similarly positioned in pairs of correct poses than in other pairs of poses ranked highly by the per-ligand scoring function (**Fig. 2C**).

These results suggest that the similarity of correct poses is not adequately captured by a state-of-the-art per-ligand scoring function. In further support of this point, we also calculated probability distributions of similarity between the poses that the per-ligand scoring function ranks first for each ligand (**Supplementary Fig. 2**). We found that these distributions also exhibited lower similarity than the corresponding native distributions.

### Derivation of a statistical potential for sets of binding poses

We used the similarity distributions described in the previous section to derive a statistical potential that—instead of acting on features of a single pose, as in previous docking software— acts on a set of hypothesized poses, one for each ligand known to bind the target protein.

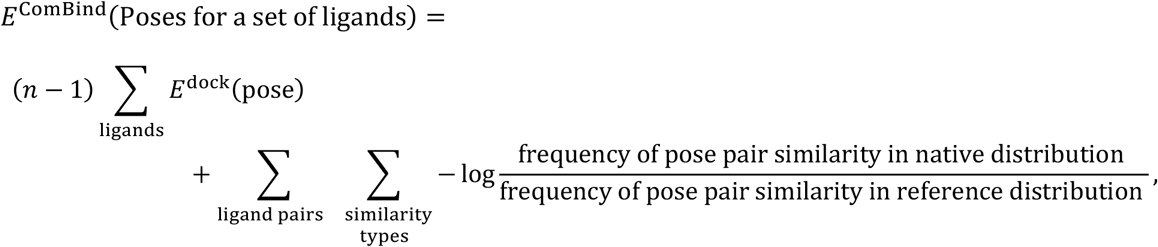

where *n* is the total number of ligands.

The first component evaluates the energetic favorability of each ligand’s pose individually using a per-ligand scoring function *E*^dock^ (e.g., a scoring function used in Glide or another docking software package). The summation is over ligands known to bind the target protein, with “pose” referring to the hypothesized pose for each ligand.

The second component rewards sets of poses with a degree of similarity that is more often observed in correct poses than in other poses ranked highly by the per-ligand scoring function. Here the outer summation is over pairs of distinct ligands known to bind the target protein, and the inner summation is over the similarity measures shown in **Fig. 2B and 2C**: hydrogen bond similarity, salt bridge similarity, hydrophobic contact similarity, and substructure similarity. “Pose pair similarity” refers to the calculated similarity value of the given type for the hypothesized poses of the given ligand pair. The “native distribution” and “reference distribution” for each similarity type are determined as described above. The resulting negative log likelihood ratios have the mathematical properties of an energy, namely that an additive decrease in energy corresponds to a multiplicative increase in likelihood ratio, allowing for straightforward integration with standard per-ligand docking scores, which are typically in units of energy (**Supplementary Fig. 3)**. For pairs of ligands that do not share a substructure at least half the size of the smaller ligand, the substructure similarity term is not included in the summation.

The second component acts as a correction to the first. If the per-ligand scoring function were perfect, in the sense that it could perfectly distinguish correct poses from incorrect ones, the terms in the second component would consistently assume values of zero. Because per-ligand scoring functions remain imperfect—and, in particular, tend to underpredict the likelihood that a set of ligands will adopt similar poses—the second component typically assumes non-zero values.

### Structure prediction informed by non-structural data

The ComBind pose prediction method identifies a set of binding poses—one for each of a set of ligands known to bind the target protein—that minimizes the ComBind potential. More specifically, given a target protein and a query ligand whose binding pose we wish to predict, we proceed as follows:

1. Compile a set of other ligands known to bind the target protein (e.g., from a public database such as ChEMBL or from ligands tested as part of a drug discovery project). We refer to these as helper ligands.
2. Dock the query ligand and each helper ligand individually to the target protein (with a per-ligand docking software package), generating many candidate poses and associated docking scores for each ligand.
3. Determine the set of poses—one per ligand—that minimizes the ComBind potential. We use an expectation-maximization algorithm for this purpose (see Methods).

As an illustrative example, we apply ComBind to predict binding poses for ligands at the β_1_-adrenergic receptor (β_1_AR), the primary target of the beta blocker drugs that are widely used to treat heart attack, heart failure, and hypertension. We selected 11 diverse ligands known to bind β_1_AR, including both beta blockers and beta agonists. We docked 11 distinct ligands to a crystallographic β_1_AR structure using Glide, producing up to 100 candidate poses for each ligand. We then solved for a set of poses—one per ligand—that minimizes the ComBind potential (**Fig. 3A**). The crystallographic β_1_AR structure used for docking was determined in complex with a ligand distinct from any of the 11 docked ligands. Crystallographic ligand poses were not used in any way by Glide or ComBind.

**Figure 3:**
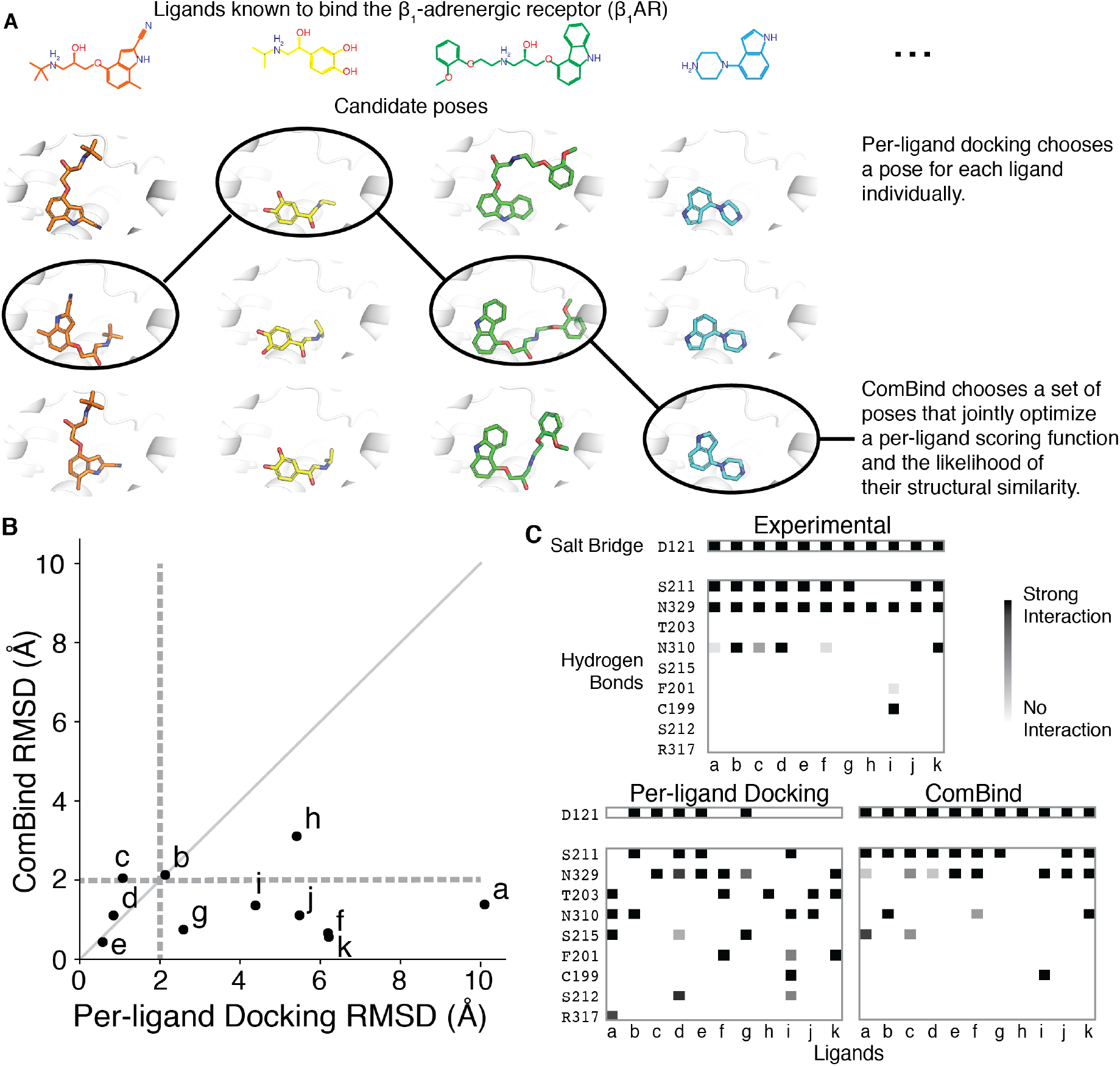
ComBind discovers and rewards key interactions shared by distinct ligands. (A) Whereas per-ligand docking considers each ligand individually, ComBind jointly selects poses for all ligands, optimizing for poses that are individually favorable according to a per-ligand scoring function and together form a coherent set of protein–ligand interactions. (B) We tested the ability of ComBind to predict the poses of 11 ligands that bind β_1_AR. Each dot corresponds to a single ligand, with the dot’s position indicating the error in the predicted pose (RMSD from the experimentally determined pose) for ComBind and for state-of-the art per-ligand docking software (Glide). A pose is considered correct if its RMSD is ≤ 2.0 Å (dashed lines). ComBind predicts a substantially more accurate pose than Glide for 7 of the 11 ligands. (C) The set of residues with which each ligand forms salt bridges or hydrogen bonds when positioned in its experimentally determined pose (top), the pose predicted by per-ligand docking (left), and the pose predicted by ComBind (right).

Glide’s top-ranked pose was correct for 4 of 11 ligands, whereas the pose selected by ComBind was correct for 10 of 11 ligands (**Fig. 3B).** In ComBind’s selected poses—as in experimentally determined poses—most of the ligands form a salt bridge with D121 and hydrogen bonds with S211 and N329 (**Fig. 3C**). In comparison, the poses ranked most highly by Glide’s per-ligand scoring function exhibited more varied hydrogen bonds and salt bridges (**Fig. 3C**).

We emphasize that ComBind does not require that all ligands adopt similar poses or form similar interactions. ComBind correctly predicts, for example, that two of these β_1_AR ligands do not form a hydrogen bond with S211.

### ComBind outperforms a state-of-the-art method on a diverse benchmark set

We benchmarked ComBind on a set of 245 protein–ligand complexes representing all major families of drug targets. We took several steps to mimic a real-world use case. First, when predicting the binding pose of a query ligand with ComBind, we used helper ligands selected from the public ChEMBL database (16). We did not use any experimental information on poses of helper ligands; indeed, for the vast majority of helper ligands selected, poses have not been determined experimentally. Second, we never used a target protein structure determined in the presence of a ligand that shares a chemical scaffold with the query ligand, ensuring that we performed only “cross-docking” and avoided the easier but less practically relevant case of “self-docking” (see Methods). Finally, when predicting ligand binding poses at a given target protein, we omitted all structures involving that protein when constructing the distributions used to define the ComBind potential.

We evaluated two ways of choosing, from ChEMBL, helper ligands for use in ComBind: (1) a diverse set of ligands with the highest binding affinity (“high affinity”), and (2) the ligands sharing the largest substructure with the query ligand (“congeneric”). Both of these selection criteria lead to substantial performance improvements over Glide (**Fig 4, Supplementary Fig. 4**), indicating that ComBind could be applied effectively using either a diverse set of ligands identified from a high-throughput screen or a congeneric series of ligands generated during lead optimization.

**Figure 4:**
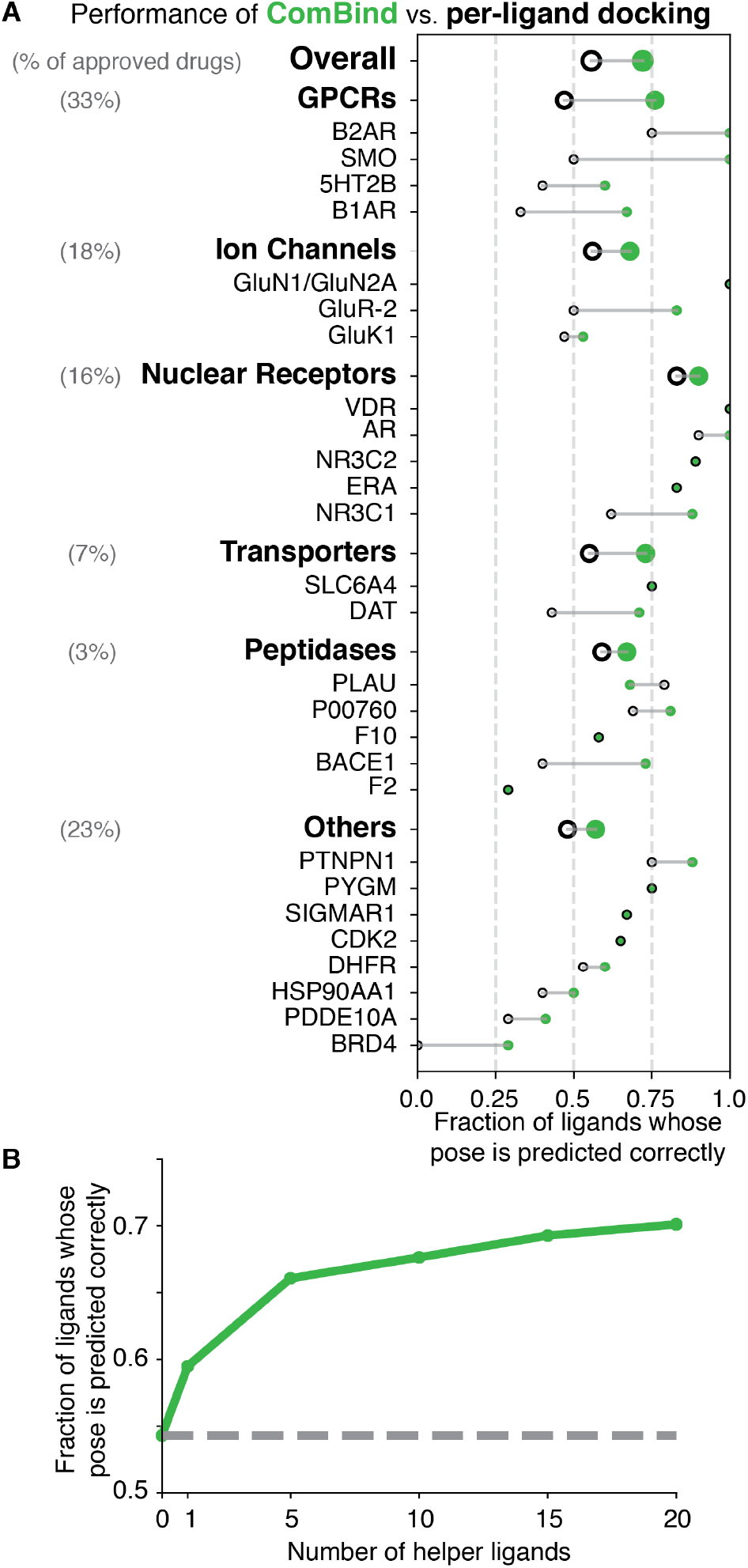
ComBind outperforms per-ligand docking on a diverse benchmark set. Performance of ComBind, as compared to a per-ligand scoring function, using helper ligands selected automatically from ChEMBL. All results are for “cross-docking” (the query ligand is docked into a structure determined in the presence of a distinct ligand). (A) Performance per target protein, target protein family (GPCRs, ion channels, etc.), and overall. Green disks and black circles indicate performance (fraction of ligands whose pose is predicted correctly) for ComBind and a state-of-the art per-ligand docking software package (Glide), respectively. (B) Performance as a function of the number of helper ligands. When using no helper ligands, ComBind is equivalent to Glide (dashed horizontal line).

ComBind’s performance improves with the use of more helper ligands (up to 20, the maximum number we tested) (**Fig. 4B, Supplementary Fig. 4B**). Interestingly, ComBind substantially outperforms Glide even when using only a single helper ligand.

In the ComBind benchmark results described below (**Fig. 4A**), we used 20 helper ligands for each query ligand, selected from ChEMBL by the high-affinity criterion. When computing overall results, we averaged across target families, weighted the performance for each family by the fraction of FDA-approved drugs targeting that family (20).

On average, ComBind selects a correct pose for 57% of all ligands and 70% of ligands for which at least one correct pose was included among the list of candidates considered—a 30% improvement over Glide in both cases (**Supplementary Table 1**). ComBind improves pose prediction performance for all target families considered. Even at the individual target level, we find that use of ComBind hardly ever degrades performance: ComBind only reduced performance for one of the 30 targets considered, and this performance reduction was minor.

Removing any of the similarity types from the ComBind potential reduced ComBind’s performance (**Supplementary Fig. 5**). In particular, both protein–ligand interaction similarity and substructure similarity contribute substantially to ComBind’s accuracy. Protein–ligand interaction similarity is the more important of the two, particularly when using a diverse set of helper ligands.

### Predicting binding poses of antipsychotics at the D_2_ dopamine receptor

To illustrate the practical application of ComBind, we predicted the binding poses of three antipsychotic drugs—pimozide, benperidol, and spiperone—at their target, the D_2_ dopamine receptor (D_2_R). Knowledge of these binding poses could aid ongoing efforts to develop antipsychotics with improved pharmacological properties, including ligands that bind selectively to D_2_R over other dopamine receptors (Butini et al., 2016; Moritz et al., 2018). Solving experimental structures of D_2_R has proven difficult, despites decades of effort (Wang et al., 2018, Yin et al., 2020). At the time we made these predictions, the only available D_2_R structure was for D_2_R bound to risperidone (22), a ligand substantially different from those whose poses we wished to predict.

We predicted binding poses for pimozide, benperidol and spiperone, as well as the tool compound mespiperone, using both Glide and ComBind (see Methods). For spiperone and mespiperone, ComBind and Glide predict similar poses. For pimozide and benperidol, however, ComBind’s predictions are different from Glide’s: a fluorobenzene ring of each compound is positioned near the top of the binding pocket by Glide and near the bottom by ComBind (Fig. 5A,B, Supplementary Fig. 6A, B).

**Figure 5:**
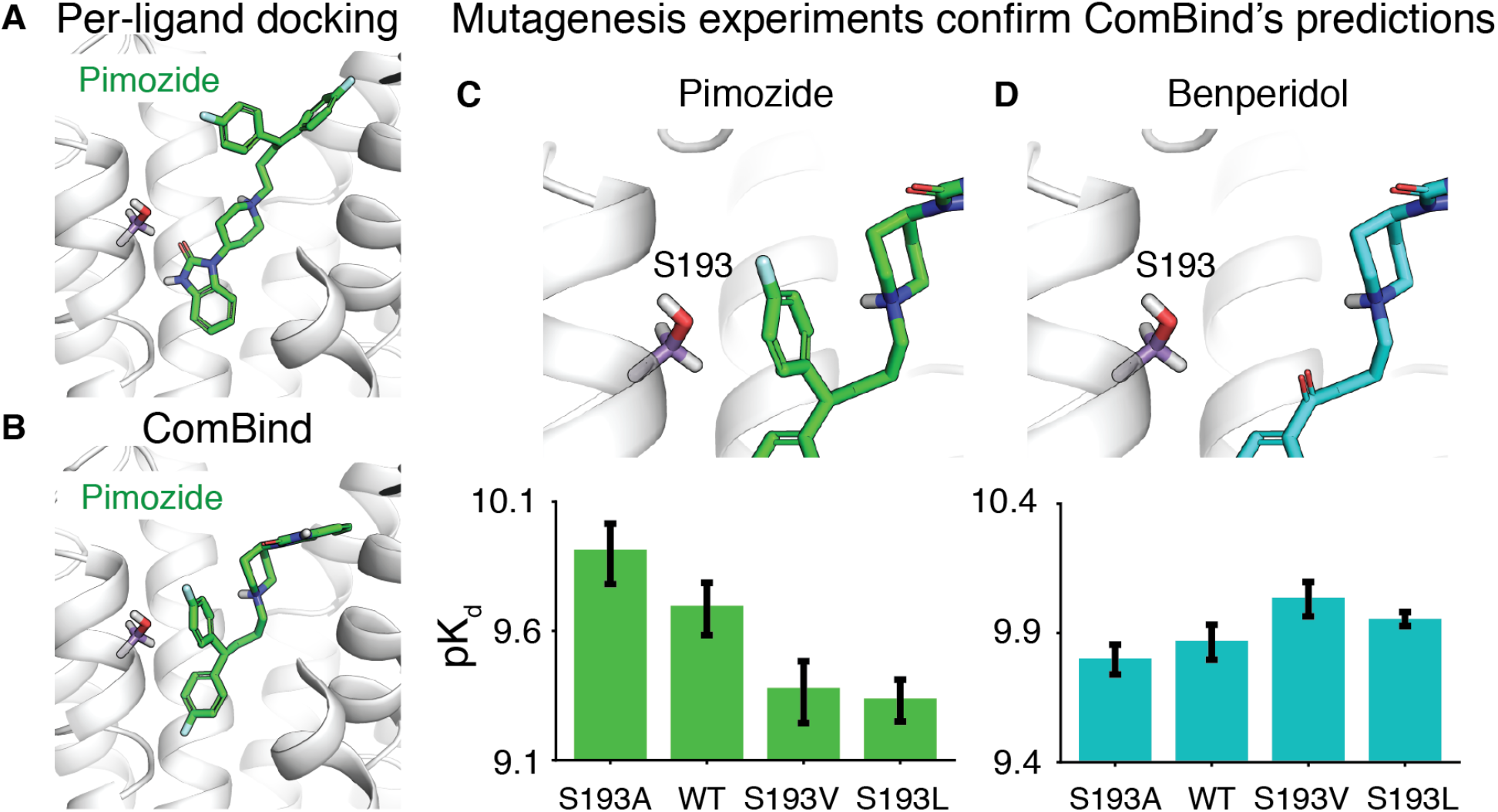
Prediction and validation of the binding poses of antipsychotics at the D_2_ dopamine receptor. Glide (A) and ComBind (B) predict very different binding poses for pimozide (and for benperidol; see **Supplementary Fig. 6**). (C) Mutagenesis experiments validate ComBind’s predictions. In ComBind’s predicted pose for pimozide, its “extra” ring is uncomfortably close to S193, such that decreasing the size of residue 193 (S193A) increases pimozide’s binding affinity and increasing the size of residue 193 (S193V and S193L) decreases pimozide’s binding affinity. WT represents the wild-type (unmutated) receptor. (D) As a control, we verified that benperidol—which lacks this “extra” ring but is otherwise identical to pimozide—does not exhibit the same trend. Error bars show standard error of the mean. See **Supplementary Fig. 6** for additional data.

To test ComBind’s predictions, we designed mutagenesis experiments. First, we tested a series of mutations of Ser193 (S193), which is positioned uncomfortably close to the second fluorobenzene ring of pimozide in ComBind’s predicted pose but not in Glide’s (Fig. 5C). Indeed, mutating S193 to a larger residue (Val or Leu) decreases pimozide’s affinity, while mutating S193 to a smaller residue (Ala) increases pimozide’s affinity. Such effects are not observed for benperidol, which is identical to pimozide except that it lacks the fluorobenzene ring that contacts S193 in pimozide (Fig. 5D). Indeed, benperidol’s affinity actually increases when S193 is mutated to a larger residue. These results are consistent with ComBind’s predicted poses but not with Glide’s: Glide predicts that pimozide and benperidol position nearly identical chemical groups in essentially identical positions near S193. Additional experiments involving mutation of residues surrounding the top and bottom of the binding pocket also support ComBind’s predictions (Supplementary Fig. 6C).

Shortly before submission of this manuscript, a haloperidol-bound D_2_R crystal structure appeared (Fan et al., 2020). Haloperidol shares a common substructure with the ligands we considered, and this substructure is positioned similarly in in the crystal structure and in ComBind’s predictions, further supporting the accuracy of these predictions.

Our results highlight a structural motif contributing to potent and selective binding to D_2_R. The antipsychotics we studied have picomolar affinity at D_2_R and bind more tightly to D_2_R than to the D3 dopamine receptor (D3R). Haloperidol, by contrast, binds with weaker (nanomolar) affinity and is not selective for D_2_R over D3R. Comparison of the binding poses reveals that the primary difference in the protein–ligand interactions is that all of the antipsychotics we studied— but not haloperidol—place a ring structure in the “extracellular vestibule,” located above the orthosteric site where dopamine binds. The extracellular vestibule has much higher sequence diversity among the different dopamine receptors than does the orthosteric site, supporting the hypothesis that ligand interactions with this region contribute to selectivity. Optimizing ligands to strengthen these interactions could lead to drugs with greater selectivity for D_2_R.

## Discussion

We have introduced a statistical potential that acts on a set of structures for different protein-ligand complexes, rather than on a single structure. We have used this potential to develop ComBind, a method that increases the accuracy of binding pose prediction by simultaneously considering the poses of multiple ligands known to bind the target.

Importantly, ComBind does not assume that all ligands considered bind in similar poses. Instead, it considers both the favorability of each individual ligand’s pose, as evaluated by a per-ligand scoring function, and the tendency of different ligands to adopt similar poses, as determined by analysis of hundreds of experimental structures. ComBind often predicts correctly that two ligands position their common scaffold differently, or that they form substantially different interactions with the binding pocket (**Supplementary Figs. 7 and 8**).

### Applicability and robustness

ComBind is broadly applicable. When benchmarking ComBind, we simply selected, from the ChEMBL database, helper ligands that bind to the same target as the query ligand, without requiring that these ligands be similar in any way. For most major drug targets, numerous binders have already been identified. Even for a completely novel target, several binders would typically be identified in the very early stages of a drug discovery project by high-throughput screening.

Binding pose prediction is important in many areas beyond drug discovery. These include the study of biological phenomena such as cellular signaling (e.g., binding of hormones and neurotransmitters), sensation (e.g., binding of odorants and flavorants), enzyme function (e.g., binding of nutrients and other metabolic substrates), and defense mechanisms (e.g., binding of toxins and antibiotics). Pose prediction is also important to understanding the effects of genetic variation on responses to both naturally occurring ligands and drugs, which is essential to personalized medicine (23). In each of these cases, multiple ligands are typically known to bind the targets of interest, and ComBind may thus be used to improve binding pose prediction.

ComBind is highly robust. This is illustrated by its accuracy in our benchmarks, which used helper ligands selected automatically according to approximate affinity values listed in the ChEMBL database. This data is noisy, not only because ligand affinities were measured by many labs using different assays, but also because the data often includes values that were inputted incorrectly (24, 25). In addition, ligands selected automatically from ChEMBL sometimes bind to completely different binding pockets on the same target.

ComBind generally produces an accurate prediction for the query ligand even when no correct candidate poses are generated for many helper ligands. **Supplementary Table 3** shows an example in which the majority of the ligands considered had no correct candidate pose; ComBind nevertheless outperformed per-ligand docking.

The per-ligand docking software used to generate and score individual ligand poses in our current implementation of ComBind treats the protein as rigid. Nevertheless, ComBind generally proves effective even when considering a set of ligands that bind diverse protein conformations. For example, the β_1_AR ligands considered in **Fig. 3** include both agonists, which bind preferentially to the protein’s active conformation, and inverse agonists, which bind preferentially to its inactive conformation (**Supplementary Table 4**).

### Relationship to previous work

ComBind builds upon several methods that combine ligand-based and physics-based information in more limited settings. Three-dimensional quantitative structure-activity relationship (3D QSAR) techniques, including field-based methods and 3D pharmacophore methods, are ligand-based approaches that consider potential 3D conformations of many ligands (26–28). These methods attempt to align ligands in three dimensions, but they do not require a structure of the target protein, and even when such a structure is available, it is typically used only in a limited way—e.g., to define excluded volume (29). 3D QSAR methods require data for a large number of binders and are generally not applied to pose prediction.

ComBind also draws inspiration from previous methods that predict binding poses of multiple known binders simultaneously. Some of these methods consider a congeneric series of ligands and require that the shared scaffold is similarly placed (30, 31). Others use either the number of similarly placed functional groups (32) or the number of shared interactions (33) between a set of docked ligands as a scoring function, assuming that the ligands adopt maximally similar poses. ComBind goes beyond these techniques in that it not only applies to any set of ligands but also provides a principled method to combine information from per-ligand docking scores with information on pose similarity across multiple ligands. This is essential to ComBind’s success in cases where ligands form substantially different interactions or position shared substructures very differently (34). Likewise, ComBind provides a principled method to combine multiple metrics of pose similarity. Indeed, ComBind’s performance drops substantially if one omits per-ligand docking scores, substructure similarity, or interaction similarity from its scoring function (**Supplementary Fig. 5**).

ComBind, like many ligand-based approaches, may also be viewed as a machine learning method (35). Most recent innovations in machine learning for drug discovery, including deep learning methods, involve complex models with many parameters that are able to fit extremely general functions. But this generality comes at a cost: such methods are typically effective only in cases where ligand data is abundant. ComBind is designed to make efficient use of any available ligand data by leveraging the physical priors encoded in structure-based approaches. This allows ComBind to improve upon the performance of a state-of-the-art docking method even when the list of other known binders is limited to a single ligand.

### Performance

Our extensive benchmarks show that ComBind outperforms a state-of-the-art per-ligand pose prediction method across all major families of drug targets. For individual targets, ComBind often substantially improves pose prediction accuracy and hardly ever degrades it. Using ComBind thus has little if any downside.

ComBind performs particularly well for certain families of targets. Its 60% improvement over Glide for GPCRs is especially noteworthy, not only because GPCRs represent the targets of one-third of all approved drugs—and a very large fraction of current drug discovery efforts—but also because experimentally determining structures of GPCRs in complex with lead compounds is often extremely difficult. Almost all experimentally determined structures of GPCRs are bound to ligands that were carefully selected for their very high affinities and residence times, often after structure determination with other ligands failed. More generally, ComBind appears to deliver an especially large improvement in pose prediction accuracy for ligands that bind to transmembrane domains of proteins, perhaps reflecting the deep, well-defined nature of these binding pockets. Experimental structure prediction tends to be particularly challenging for transmembrane proteins, highlighting ComBind’s utility.

ComBind’s performance could undoubtedly be improved further through use of curated or in-house data. In particular, a careful human curator could (1) identify ligands that can most confidently be classified as binders (e.g., based on multiple reports or on particularly reliable data sources), (2) identify ligands demonstrated to bind in the same binding pocket (e.g., by competition binding assays), and (3) remove data that was inputted incorrectly to a database. For a major drug discovery project focused on a particular target, a substantial amount of additional in-house data will often be available on ligands found to bind the target, and that data will typically have been collected in a more uniform and consistent manner than data extracted from multiple publications.

Skilled chemists can often improve the overall success rate of docking through careful manual preparation of the protein structure—for example, by diligent placement of waters or consideration of side chain rotamers. Such a procedure is subjective and was thus not employed in our performance benchmarks. In our experience, however, careful manual preparation of target structures improves ComBind’s results even more than those of per-ligand docking methods, because such preparation increases the accuracy of the helper ligand poses and thus the value of the information gleaned from them.

A variety of “flexible docking” methods have been developed that allow deformation of the target protein when sampling ligand poses (19, 36, 37). These methods have proven highly valuable in cases where the user knows in advance that protein flexibility is important to binding of the query ligand. When used as fully automated pose prediction methods without such prior information, however, flexible docking methods generally underperform rigid docking methods such as Glide, as observed in our benchmarks of the popular Induced Fit Docking method (36) (**Supplementary Table 1)** and reported previously for other flexible docking methods (37). Such methods are more likely to sample a correct pose but also more likely to sample incorrect poses that outscore correct poses. The ComBind scoring function might address this problem by more effectively selecting a correct pose from among the incorrect poses; this is a potential area for future research.

### Extensibility and future work

Because ComBind can use any per-ligand docking method for pose generation and scoring of individual ligands, it will be able to take advantage of improvements to these methods. For example, several recent methods use machine learning to fit scoring functions (38–40), and others allow for binding pocket flexibility when generating candidate poses (8, 36).

Likewise, ComBind can be used with any pairwise pose similarity metric or combination thereof. ComBind’s performance could potentially be improved by using more fine-grained interaction descriptors (41, 42) or by using similarity metrics based on field-based methods developed for virtual screening (28, 43).

The statistical potential used by ComBind is sufficiently general that the method could be extended to exploit other types of data, ranging from multiple experimental structures of the protein in complex with different ligands to effects of protein mutation on ligand binding. Likewise, future work might exploit the affinity of each known binder; we have not done so here to avoid obscuring the general applicability of our method, as the affinity estimates available in public databases are often determined by different techniques and thus difficult to compare to one another.

Beyond binding pose prediction, our work suggests rich opportunities to improve prediction of diverse ligand properties by combining physics-based and ligand-based modeling. For example, both physics-based and ligand-based approaches are currently used to predict or rank ligand binding affinities in order to enable virtual ligand screening. Physics-based approaches require the use of various approximations that introduce error, while ligand-based approaches are limited in their ability to predict affinities of ligands very different from those for which experimental data is available. A careful combination of the two, perhaps based on the ComBind scoring function, might outperform either one alone. Physical modeling allows ligand data to be used more efficiently by facilitating representation of the ligands in terms of specific interactions they form with the target, a level of abstraction where even chemically diverse ligands share features. Indeed, such approaches might prove effective even for predicting functional activity values whose physical basis is not known a priori. Further work will be necessary to explore these possibilities.

## Acknowledgements

We thank B. Kelly, N. Latorraca, A. Venkatakrishnan, and N. Sohoni for advice and guidance in the early stages of the project, all members of the Dror lab for insightful comments, and J. Javitch for providing materials for mutagenesis experiments. This work was supported by National Institutes of Health grants R01 GM127359 (to R.O.D.), R01-GM083118 (to R.K.S.) and U19-GM106990 (to R.K.S.), and by a Stanford Graduate Fellowship (to J.M.P.).

## Methods

### Assembly of data for use in learning the ComBind scoring function

#### Curation of experimental protein–ligand complex structures

In order to learn the ComBind scoring function, we curated a set of protein–ligand complex structures representing each of the major drug targets catalogued by Santos *et al.*, 2017 (**Supplementary Table 2**). This set of target proteins was chosen through a combination of manual curation and adaptation of the PDBbind refined set (44). For each target, we included up to 21 structures, each with a distinct ligand bound, selecting the structures with alphabetically lowest PDB code when more than 21 were available. Structures with duplicate ligands, mutant proteins, or no small molecule in the orthosteric site were excluded.

#### Generation of docked poses

For all of the results presented in this study, we performed “cross-docking.” Specifically, for each target, we chose the structure with the alphabetically first PDB code as the input 3D structure of the protein and then docked other ligands to this reference structure. This simulates a real-world application where only one structure of the target protein is available, and the user wants to predict poses for ligands not present in that structure.

To prepare protein structures for use in docking, we first prepared structures using the Schrodinger suite. All waters were removed, the tautomeric state of the ligand present in the experimentally determined structure was assigned using Epik at pH 7.0 +/– 2.0, hydrogen bonds were optimized, and energy minimization was performed with non-hydrogen atoms constrained to an RMSD of less than 0.3 Å from the initial structure. The ligand was then removed.

For ligands to be docked, the tautomeric state was assigned using Epik tool at target pH 7.0 +/– 2.0. The single most favorable state was considered for docking. Torsion angles were randomized before docking.

Ligands were docked using default Glide SP settings except that “Enhanced Sampling” was set to 4, quadrupling the number of ligand conformers considered. For each ligand, we produced up to the 100 most highly ranked poses (for some ligands fewer than 100 poses passed Glide’s internal filters). We also considered using Glide XP but found that Glide XP produced a correct candidate pose substantially less often than Glide SP (**Supplementary Table 1**). Glide XP and SP performed similarly in terms of how frequently the top-ranked pose is correct. Additionally, we considered using Induced Fit Docking (IFD). While IFD produced at least one correct candidate pose more often than Glide SP, the performance in terms of how often the top-ranked pose is correct was worse.

#### Determining the quality of docked poses

The accuracy of each pose was quantified by the non-hydrogen-atom RMSD from the experimentally determined pose. To compute the RMSD, each complex was aligned to the structure used for docking based on non-hydrogen-atoms within 15 Å of the ligand, and the RMSD was then computed between the docked pose and the same ligand’s pose in the aligned complex. We denote poses at most 2.0 Å RMSD from their aligned experimentally determined pose as being “near-native” or “correct.”

### Quantifying the similarity of binding poses for distinct ligands

#### Protein–ligand interaction similarity

Three interaction types were considered: hydrogen bonds, salt bridges, and hydrophobic contacts. We designed quantitative measures to assess the presence of these interactions between the ligand and a given protein residue (**Supplementary Table 5**). The hydrogen bond and salt bridge interaction measures were designed to give a value of 1 for interactions meeting established criteria (17). A soft boundary was added to give borderline cases values between 0 and 1, in order to prevent discontinuities. The hydrophobic contact measure approximates the hydrophobic surface contact area by considering the number of protein–ligand atom pairs in contact with each other. Again, a soft boundary (in this case, between an atom pair being or not being in contact) was used to prevent very similar poses from leading to very different values. We denote the interaction value for interaction type *k*, for pose *ℓ_i_* of ligand *i*, with protein residue *r* as 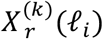.

Interaction similarities for a pair of poses (for two different ligands bound to the same target protein) were computed separately for each interaction type. The interactions made between the ligand and each residue of the target protein residue were tabulated and then the similarity between the resulting lists for each pose was measured by the Tanimoto coefficient (21). The Tanimoto coefficient was modified by the addition of pseudo counts, which serve to make the metric well defined if neither ligand forms a particular type of interaction and to reward poses that share larger numbers of interactions in absolute terms. We define the interaction similarity, for interaction type *k* between a pair of poses *ℓ_i_, ℓ_j_* (for ligands *i* and *j, respectively*), as

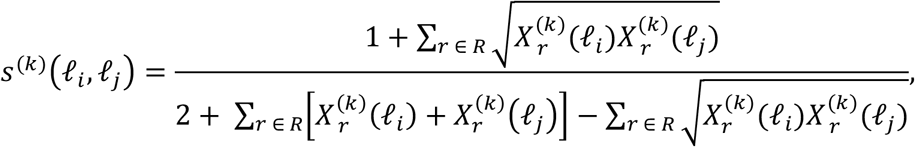

where *R* is the set of all protein residues.

When computing hydrogen bond similarity, a case where a given protein residue acts as a hydrogen bond donor for one ligand and a hydrogen bond acceptor for another ligand is not considered a shared interaction.

#### Substructure similarity

To compute the substructure similarity for a pair of candidate poses, the maximum common substructure of the two ligands is identified using Canvas (Schrodinger LLC) and then mapped onto each candidate pose. Finally, the RMSD between these two sets of atoms is computed and used as the measure of substructure similarity. We defined custom atom and bond types for computation of the common scaffold (**Supplementary Table 6**). Scaffold similarity is not considered for pairs of ligands with a maximum common substructure of less than half the size of the smaller ligand. Hydrogen atoms were not included in the substructure nor when determining the total number of atoms in each ligand.

#### Computation of similarity statistics

Using the set of protein–ligand complex structures described above, we characterized the extent to which distinct ligands binding a common target adopt similar poses, as quantified by the interaction and substructure similarity metrics described above. (We note that the three ion channel targets were not included in these statistics because they were added after the rest of our study had been completed.)

When computing these statistics, we docked the ligands using Glide and then identified poses that are near-native among the candidate poses ranked in the top 100 by Glide. We used these docked poses, as opposed to the experimentally determined pose, in order to ensure that the statistics will be applicable to the scoring of candidate poses generated by Glide. We computed the empirical distribution of each similarity type across all pairs of near-native poses using a Gaussian kernel density estimate with standard deviation of 0.03 for interaction similarities and 0.18 for substructure similarities. To reduce bias near the boundaries, we applied reflected boundary conditions(45). We capped substructure similarities at 6 Å (that is, substructure similarities greater than 6 Å were set to 6 Å), as the sparsity of near-native pose pairs for higher values led to overly rough distributions. We denote the similarity distribution over near-native poses for interaction type *k* as *f_k_*(*x*; Native).

We computed equivalent similarity distributions using all pairs of candidate poses produced by Glide, regardless of whether they are near-native. We denote the resulting distributions as *f_k_*(*x*; Reference).

To combine the distributions for the four similarity types into a single joint distribution, we assume that the interaction types are conditionally independent and express the joint distribution as a product of the distributions for each interaction type. That is:

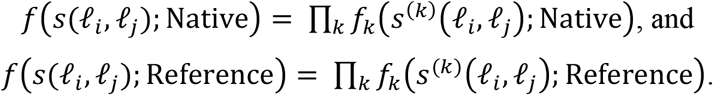

where *s*(*ℓ_i_,ℓ_j_*) is the vector of *s*^(*k*)^, (*ℓ_i_, ℓ_j_*)’s for each similarity type *k*.

### Description of the ComBind method

#### The ComBind score

We describe a hypothesized set of binding poses of a set of *n* ligands as *L* = *ℓ*_1_, *ℓ*_2_,…, *ℓ_n_*, where *ℓ_i_* specifies the hypothesized pose for ligand *i*.

Per-ligand scoring functions, which consider each ligand independently, would determine an optimal set of poses 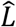 by choosing the binding pose with minimum docking score for each ligand or, equivalently, by minimizing

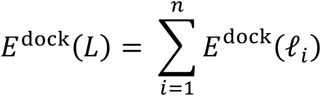

where *E*^dock^(*ℓ_i_*) is the output of a per-ligand scoring function (such as that reported by Glide) for pose *ℓ_i_* of ligand *i*.

In our method, we add pairwise terms that tend to favor sets of similar poses:

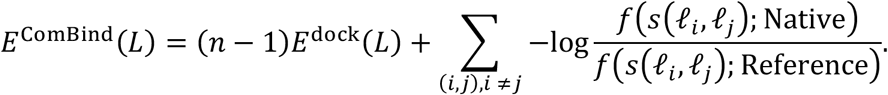

Intuitively, these pairwise terms reward pose pairs with similarity values more often observed in near-native (correct) pose pairs than in reference pose pairs (i.e., pose pairs chosen at random from among all candidates). The idea of comparing the distribution of features in correct solutions to the distribution in all possible solutions has been used in statistical potentials for biomolecular structure prediction(46–48) and in the naïve Bayes machine learning model(49). We weight the docking scores by the number of ligands minus 1, in order to hold the relative contribution of singleton and pairwise terms constant for different numbers of helper ligands.

Consistent with their reported units of kcal/mol, we find that Glide scores have the mathematical properties of an energy; namely, the negative log likelihood ratio of a pose being near-native is linear in its Glide score (**Supplementary Fig. 3**). By construction, the pairwise terms we introduce in this study also have this property. This congruence implies that these singleton and pairwise terms can be additively combined (as this is the equivalent of multiplying likelihood ratios).

In general, it could be that the per-ligand docking scores need to be scaled by a constant factor in order to be consistent with the pairwise terms. For example, if the docking scores were on average 10 times the negative log likelihood ratio of a pose being near-native, they would need to be scaled by 1/10. This constant factor can be identified by performing logistic regression with the docking scores as features and whether each pose is near-native as the response. For Glide scores, the appropriate constant is close to 1 (1/0.9 = 1.1) (**Supplementary Fig. 3**), and we chose to set it to 1 for simplicity.

#### Optimization procedure

We use coordinate descent to compute a set of poses that minimizes the ComBind score. At first, *L* is randomly initialized. *L* is then iteratively improved by iterating through the ligands, in a random order, and updating the selected ligand’s pose to the argument minimum of £ComBmd(£) assuming that the other poses in *L* are correct. This procedure is repeated until no more updates can be made. Each update can be computed efficiently because it depends only on the partial contribution of the selected ligand’s pose to the ComBind score:

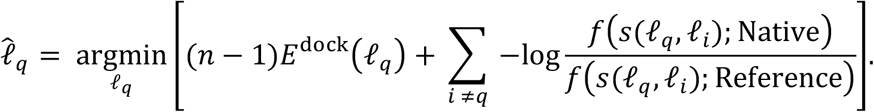

In order to account for the non-convex nature of the ComBind score, we repeat this algorithm from 500 initial configurations, explicitly including the initial configuration corresponding to the generic scoring function predictions at least once and return the best scoring configuration. Empirically this procedure converges to the same result over multiple runs.

#### Benchmarking

We evaluated the performance of ComBind on the 30 target proteins listed in **Supplementary Table 2.** We only considered ligands that have less than 50% scaffold overlap with the ligand that was originally present in the experimental structure used for docking. We found that ligands with higher scaffold overlap were substantially easier to dock, likely due to the binding pocket being well shaped to accommodate the similar ligand (**Supplementary Table 1**). Additionally, we only consider ligands for which there is at least one correct candidate pose, since only in these cases is it possible for either ComBind or Glide to make a correct prediction. Importantly, this subsetting was only done for the query ligands, not the helper ligands downloaded from ChEMBL described below.

For each of the 245 unique ligands meeting these criteria, we identified other ligands known to bind the respective target protein from the ChEMBL database and then used ComBind to jointly predict their binding poses. Importantly, when evaluating the performance of our method on a particular target protein, we excluded the data for that target protein from the similarity statistics.

#### Selection of helper ligands

For all targets, we downloaded K_i_ or IC_50_ data (whichever was more numerous) from ChEMBL (16). We removed ligands that did not meet the following criteria: a ChEMBL confidence score of 9 (the highest value), molecular weight < 800 Da, and K_i_ or IC_50_ < 1 μM. Ligand structures were generated from the SMILES strings provided by ChEMBL.

We benchmarked two criteria for selecting which ChEMBL ligands to use as helper ligands for each query ligand: (1) the highest affinity binders that do not share a chemical scaffold, and (2) the ligands that share the largest chemical substructure with the query ligand. To define the size of the common substructure, we used the same maximum common substructure definition as that used to compute substructure similarity. For selection method (1), we added helper ligands in order of affinity, not adding a ligand if it has greater than 80% substructure overlap with any ligand already in the selected set of helpers.

The benchmarking results presented in the figures were obtained using the following ligand selection criteria and number of helper ligands: **Fig. 4A** and **Supplementary Fig. 5A**: 20 helper ligands selected using criterion (1); **Fig. 4B**: the indicated number of ligands selected using criterion (1); **Supplementary Fig. 4A** and **Supplementary Fig. 5B**: 20 helper ligands selected using criterion (2); and **Supplementary Fig. 4B**: the indicated number of ligands selected using criterion (2). For a handful of targets, fewer than 20 helper ligands were available meeting our criteria. In these cases, we used the minimum of the indicated number of ligands and the number of available ligands. Targets with only one ligand are omitted from **Fig. 4A** and **Supplementary Fig. 4A**.

#### Performance evaluation

We developed an overall performance metric to represent the expected performance in drug development campaigns. For each protein family, we computed the average performance, then weighted each by the fraction of FDA-approved drugs targeting the protein family, as reported in Santos *et al*., 2017.

### Prediction of binding poses of antipsychotics at the D_2_R

#### Execution of the ComBind method

We predicted binding poses for the typical antipsychotics spiperone, mespiperone, benperidol, and pimozide at the D_2_ dopamine receptor (D_2_R). We prepared the ligands using the Schrodinger ligprep tool, considering the unprotonated tautomer and both inversions of the protonated tautomer. The same docking protocol was used as described above, except that the top 300 poses were considered by ComBind, in order to account for the use of the 3 tautomeric states of the ligand.

#### D_2_ Dopamine receptor mutagenesis

Wild type (wt) human D_2_R in pcDNA3.1 was kindly provided by the laboratory of Jonathan Javitch (Columbia University, New York, NY). Mutations were introduced through of a modified QuikChange (Stratagene, La Jolla, CA) mutagenesis protocol using the following primers V91F: 5’-GGTCATGCCCTGGTTTGTCTACCTGG-3’, S193A: 5’CGTGGTCTACGCCTCCATCGTCTCC-3’, S193V: 5’-CGTGGTCTACGTCTCCATCGTCTCC-3’, S193L: 5’-CGTGGTCTACCTCTCCATCGTCTCC-3’, W100L: 5’-GGTAGGTGAGTTGAAATTCAGCAGG-3’, C118M: 5’-GGACGTCATGATGATGACGGCGAGC-3’, W386F: 5’-CGTGTTCATCATCTGCTTTCTGCCCTTCTTC-3’, F389L: 5’-GCTGGCTGCCCTTATTCATCACACACATCC-3’.

#### Membrane preparation and radioligand binding

Membranes were isolated from HEK293T cells transiently transfected with D_2_R(wt) or D_2_R-mutants. Briefly, cells were harvested 48 hr post-transfection (with Lipofectamine 2000), rinsed with PBS, lifted with harvesting buffer (0.68 mM EDTA, 150 mM NaCl, 20 mM HEPES, pH 7.4), and centrifuged at 200 x g for 3 min. The cells were resuspended in ice cold homogenizing buffer (10 mM HEPES, pH 7.4, 100 mM NaCl, 0.5 mM EGTA), homogenized using a Tissue Tearer (BioSpec, Bartlesville, OK) for 30 sec, and centrifuged at 20,000 x g for 20 min. Membranes were resuspended in Binding Buffer (20 mM HEPES, pH 7.4, 100 mM NaCl) using a Dounce glass homogenizer, flash frozen in liquid N_2_ and stored at –80°C.

For saturation binding assays, cell membranes (0.6–20 μg per well, depending on the mutant) were incubated for 1.5 hr at 30°C with [^3^H]-spiperone (Perkin Elmer, Waltham, MA) (0.02–12 nM, depending on the Kd of the D_2_R mutant) in Binding Buffer containing 0.001% BSA, 1 mM ascorbic acid, and 100 nM GDP with or without 20 μM (+)butaclamol (to determine non-specific binding). For competition binding assays, cell membranes (0.6–20 μg, depending on the D_2_R mutant) were incubated for 1.5 h at 30°C with [^3^H]-spiperone (0.05–0.6 nM, depending on the Kd of the D_2_R mutant) in Binding Buffer containing 0.001% BSA, 1 mM ascorbic acid, 100 nM GDP and 0–0.1 nM test compound (purchased from Millipore-Sigma, St Louis, MO), or 20 μM (+)butaclamol (to determine non-specific binding). Sample membranes were harvested by vacuum filtration on 96-well GF/C filter plates, washed with ice cold binding buffer to remove unbound radioligand, and allowed to dry before adding Microscint 0 (Perkin Elmer, Waltham, MA) for counting in a Top Count Scintillation Counter (Perkin Elmer/Packard, Waltham, MA). Data were fit to a one site binding curve to determine Kd for [^3^H]-spiperone saturations, or to a one-site competition binding curve to calculate Ki of test compounds using Prism (GraphPad, San Diego CA).

## Supplementary Figures

**Supplementary Figure 1:**
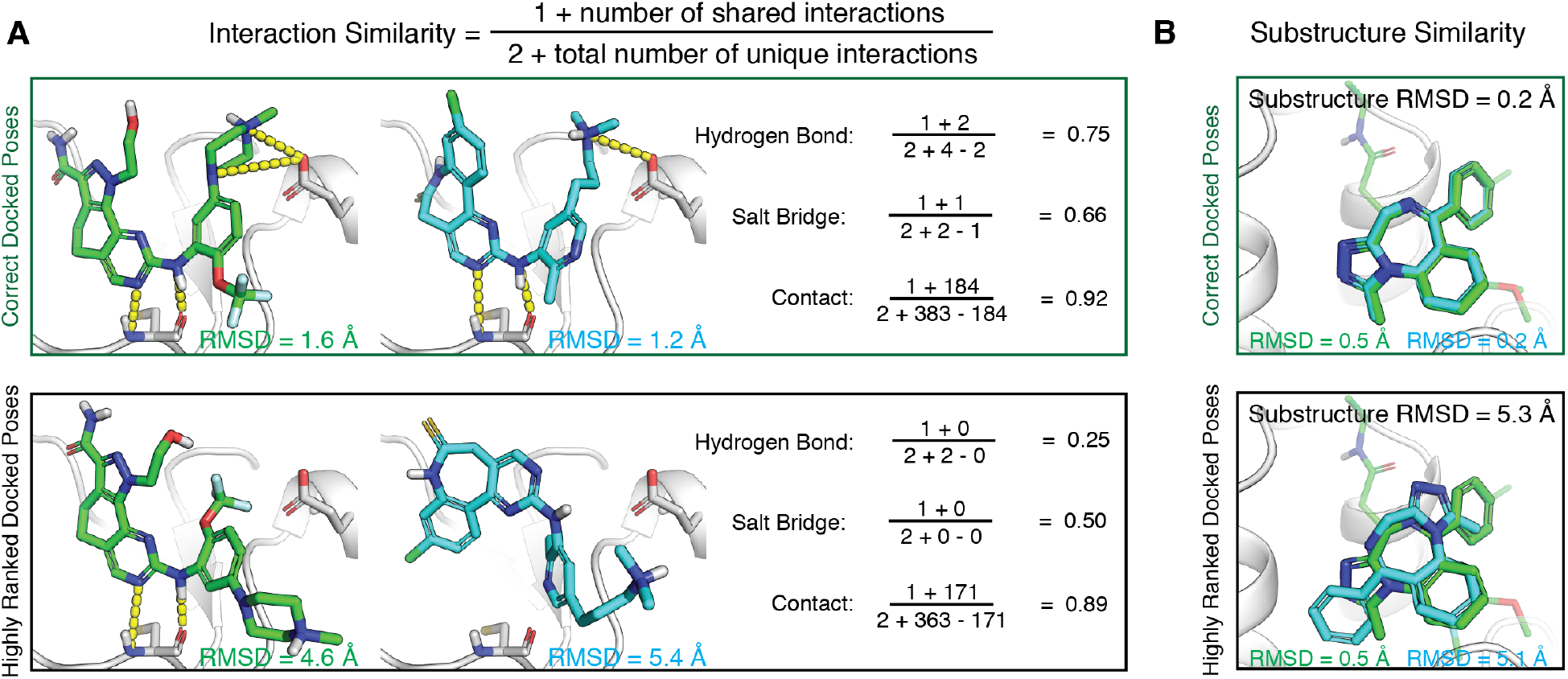
Examples of interaction similarity and substructure similarity computation. (A) Comparison of interactions formed by two ligands bound to *PLK1*, for a pair of correct poses (top) and randomly chosen poses (bottom). (B) Overlays of two ligands that share a common substructure bound to *BRD4* for correct docked poses (top) and randomly chosen highly ranked docked poses (bottom).

**Supplementary Figure 2:**
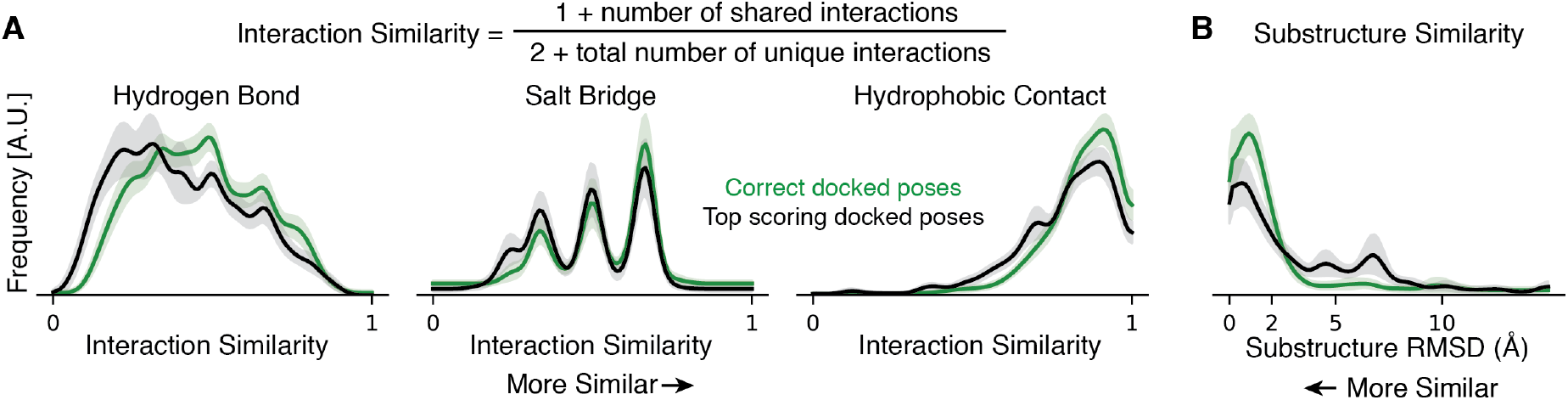
A state-of-the-art per-ligand scoring function (Glide) underestimates the similarity of binding poses of different ligands binding to the same target protein. (A) and (B) are identical to Fig. 2C and D, respectively, except that the black curves in this figure are computed using only the pose ranked first by Glide for each ligand.

**Supplementary Figure 3:**
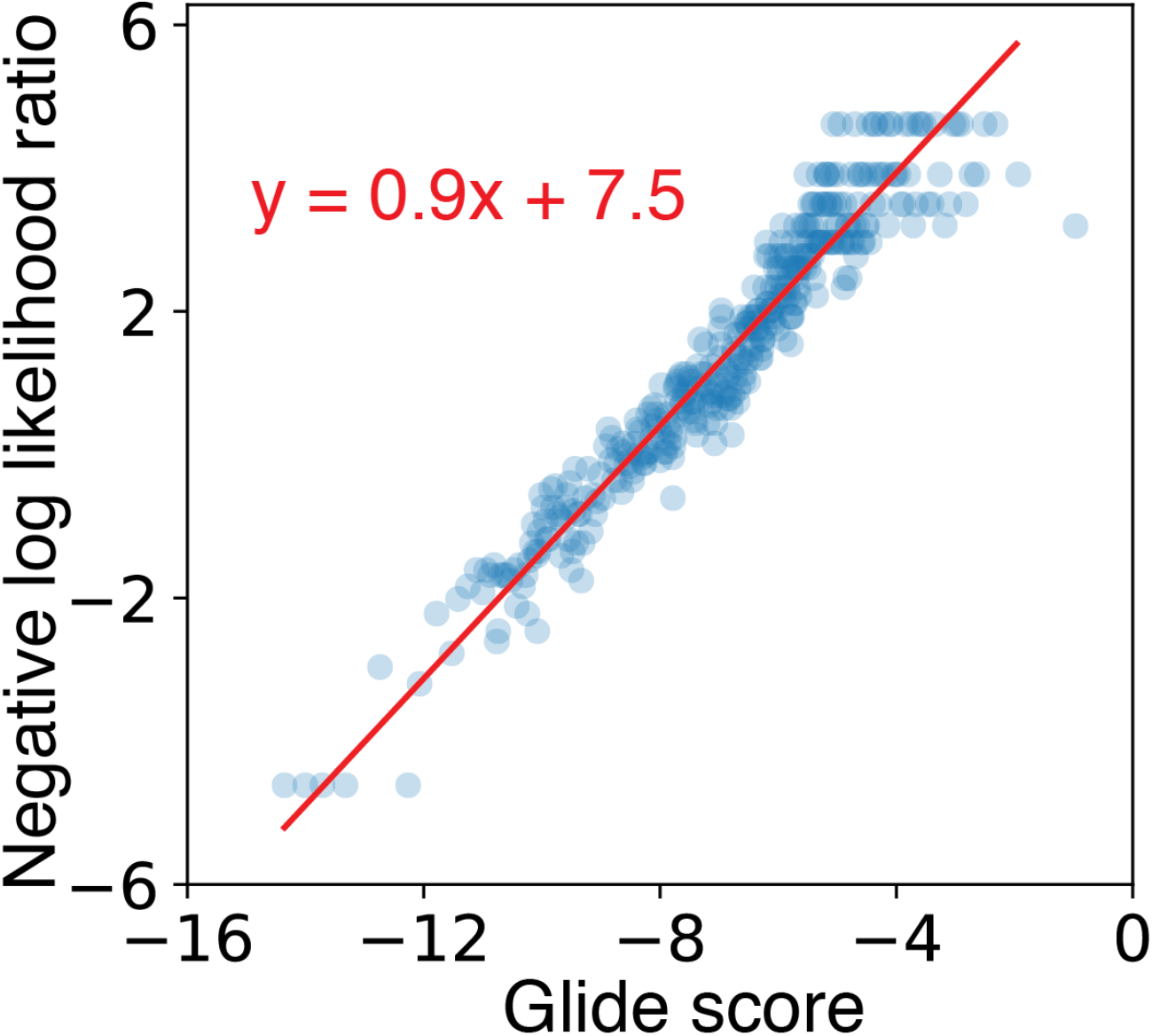
The output of Glide’s per-ligand scoring functions is in units of energy similar to those of ComBind’s pairwise potential. A quantile plot showing the relationship between Glide scores and the negative log likelihood ratio of a pose being correct. For each of the docked poses of each ligand in our benchmark set, we computed the Glide score and determined whether the pose was correct. We split all of the resulting data into quantiles based on Glide scores, with each quantile containing 100 poses. Each point in the plot represents the mean Glide score and negative log likelihood ratio for a given quantile. The red line shows the best-fit linear relationship between these two quantities as determined by logistic regression.

**Supplementary Figure 4:**
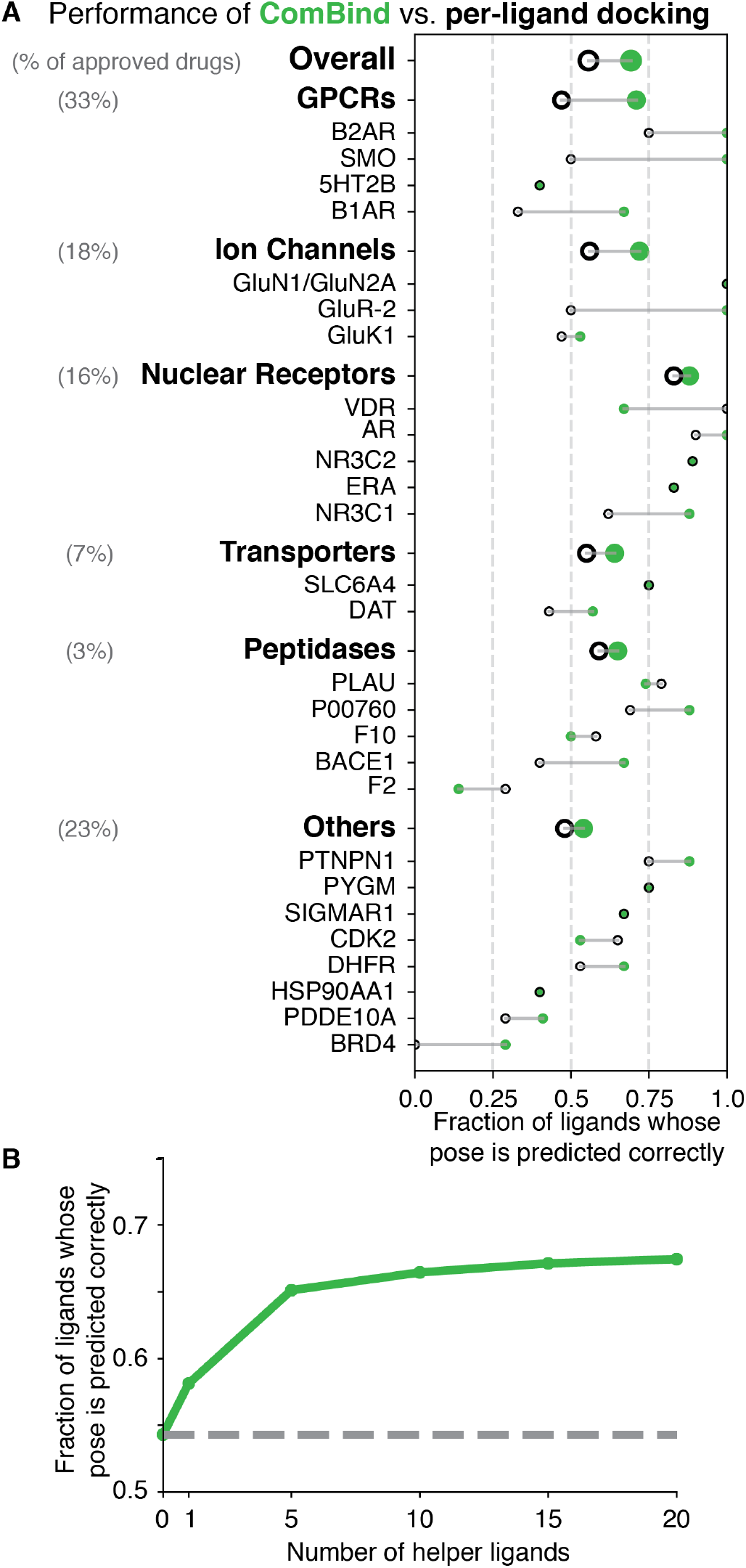
ComBind performance using a congeneric series of ligands. This figure corresponds to **Fig. 4**, but with helper ligands selected from ChEMBL ligands according to the “congeneric” criterion (i.e., ligands that share the greatest common substructure with the query) instead of the “high affinity” criterion.

**Supplementary Figure 5:**
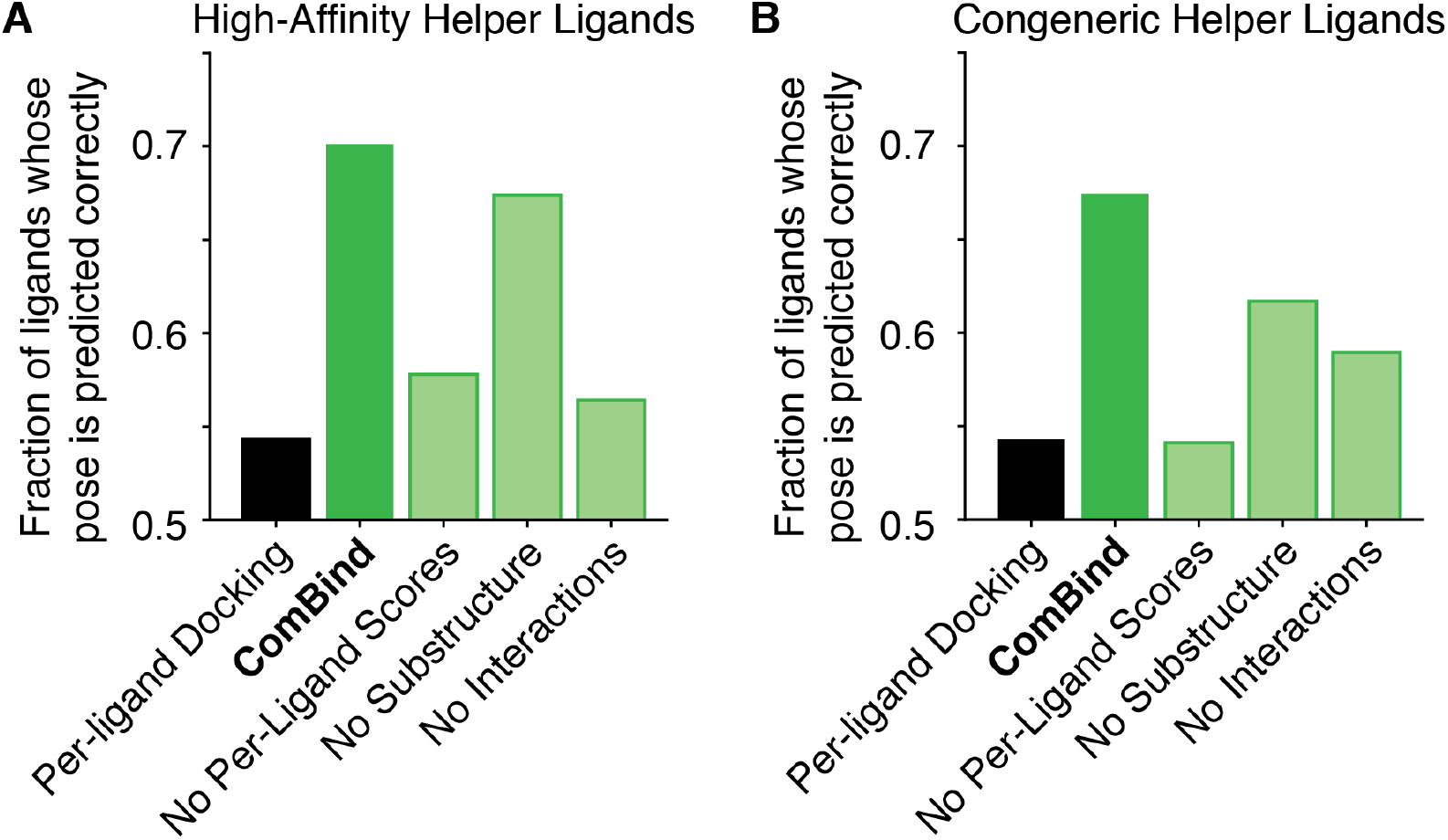
Importance of components of the ComBind scoring function. Performance using various components of the ComBind scoring function when using helper ligands chosen by either the high-affinity (A) or congeneric (B) ChEMBL ligand selection criterion. ComBind uses per-ligand docking scores, similarity scores based on interactions, and similarity scores based on relative positions of shared substructures. “Per-ligand docking” (Glide) omits all similarity scores. The remaining bars (“No Per-Ligand Scores,” “No Substructure,” and “No Interactions”) show the effects of omitting per-ligand scores, substructure position similarity scores, and protein–ligand interaction similarity scores, respectively, from the ComBind potential.

**Supplementary Figure 6:**
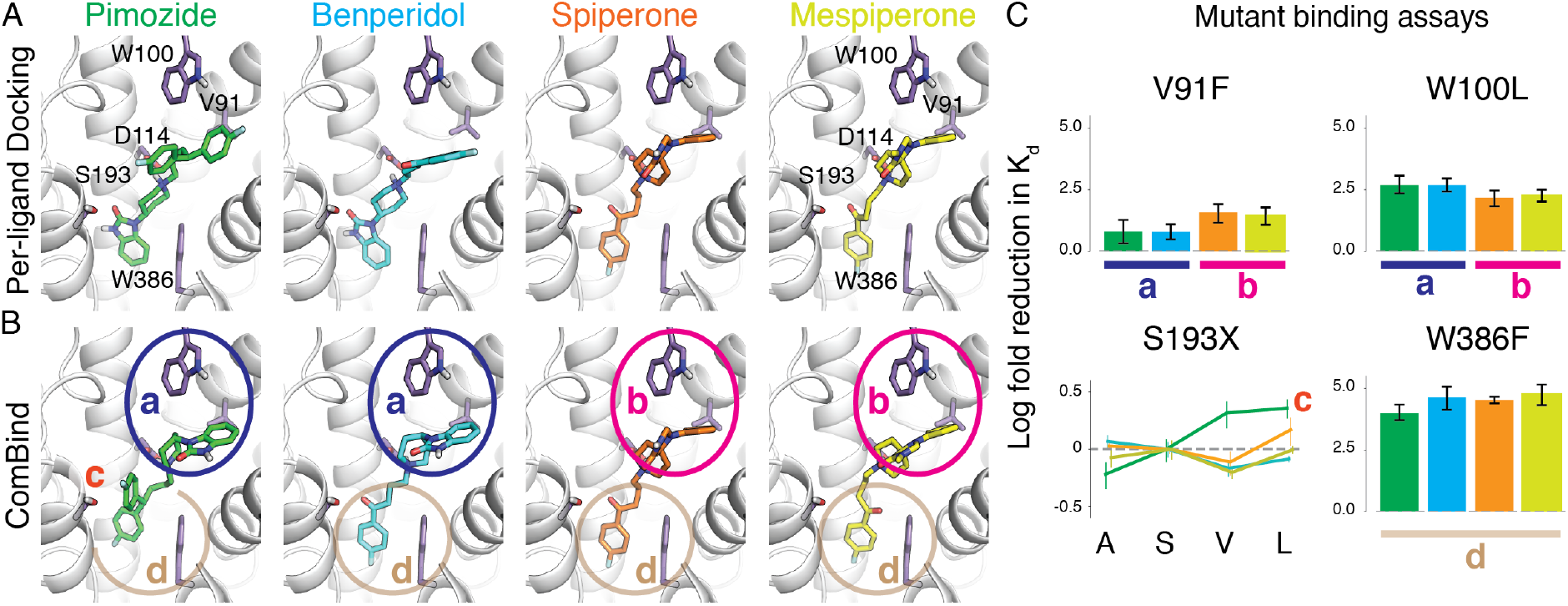
Prediction and validation of the binding poses of antipsychotics at the D_2_ dopamine receptor—additional data. (A) Binding poses of pimozide, benperidol, spiperone, and mespiperone as predicted by Glide. (B) Binding poses of the same ligands, as predicted by ComBind. (C) Results of mutagenesis studies designed to test ComBind’s binding pose predictions. Ligands are color-coded as in panel A. Error bars show standard error of the mean. S193 was mutated to A, S, V and L; these results are discussed in the main text. Unlike Glide, ComBind predicts that all four ligands will position a fluorobenzene ring at the bottom of the binding pocket, packing favorably against Trp386 (W386). Indeed, mutating W386 to a smaller residue (Phe) reduced affinity to a similar extent for all of the ligands, with a slightly smaller effect for pimozide, which packs less tightly against W386 according to ComBind’s prediction. At the top of the ligand binding pocket, near Val91 (V91) and Trp100 (W100), ComBind predicts that the pimozide and benperidol will place identical functional groups that differ somewhat from those of spiperone and mespiperone. Indeed, mutation of these residues affects pimozide and benperidol slightly differently from spiperone and mespiperone.

**Supplementary Figure 7:**
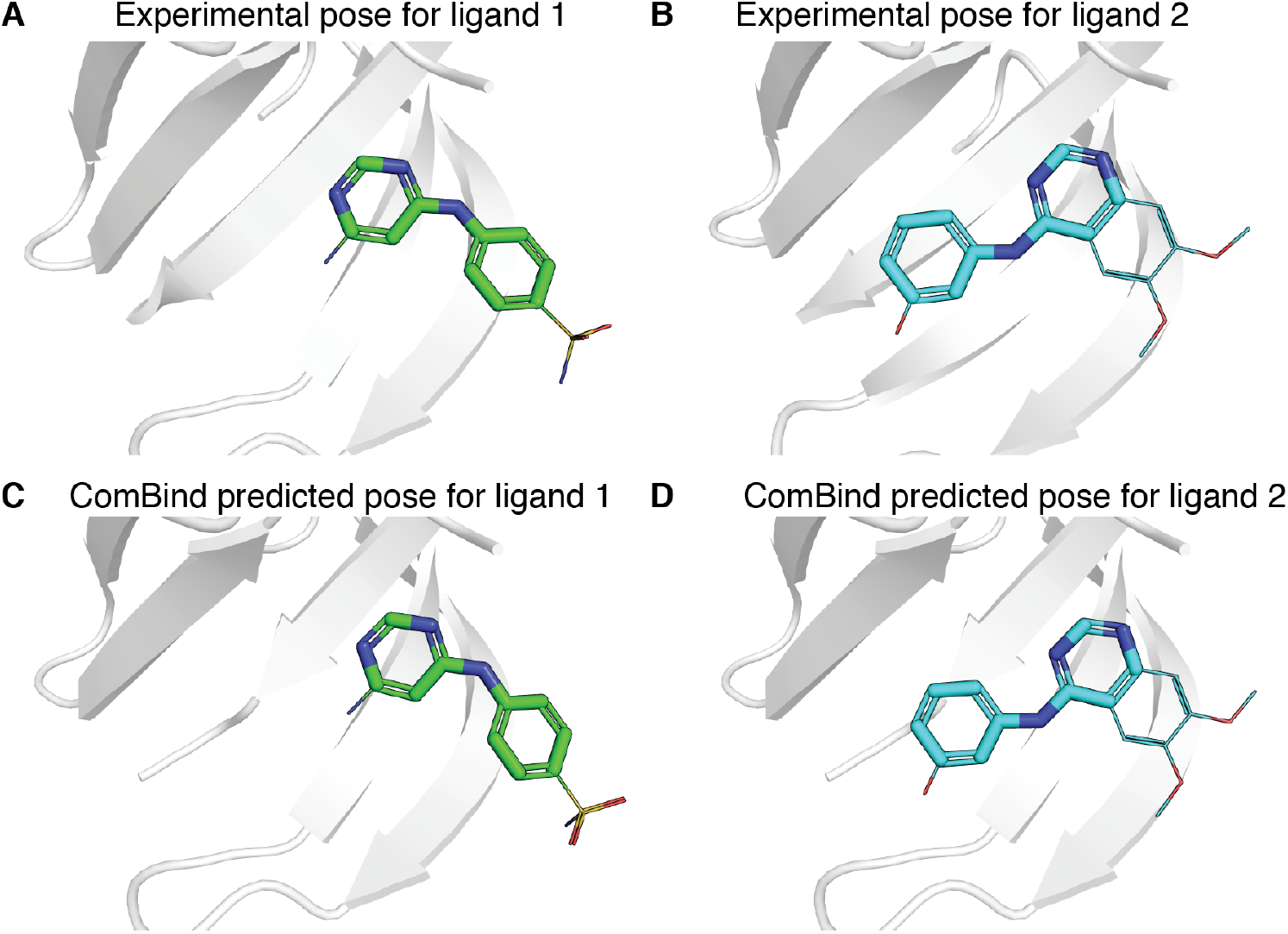
Example of a case where ComBind correctly predicts that a shared chemical scaffold is placed differently for different ligands. We show two ligands that bind the kinase CDK2. These ligands share a common scaffold but adopt significantly different binding poses. In A and B, we show their experimentally determined poses (PDB: 1JSV and PDB: 1DI8, respectively). In C and D, we show the poses predicted by ComBind for the two ligands. The shared scaffold is shown in the thicker sticks and parts of the ligands that differ are shown in the thinner lines.

**Supplementary Figure 8:**
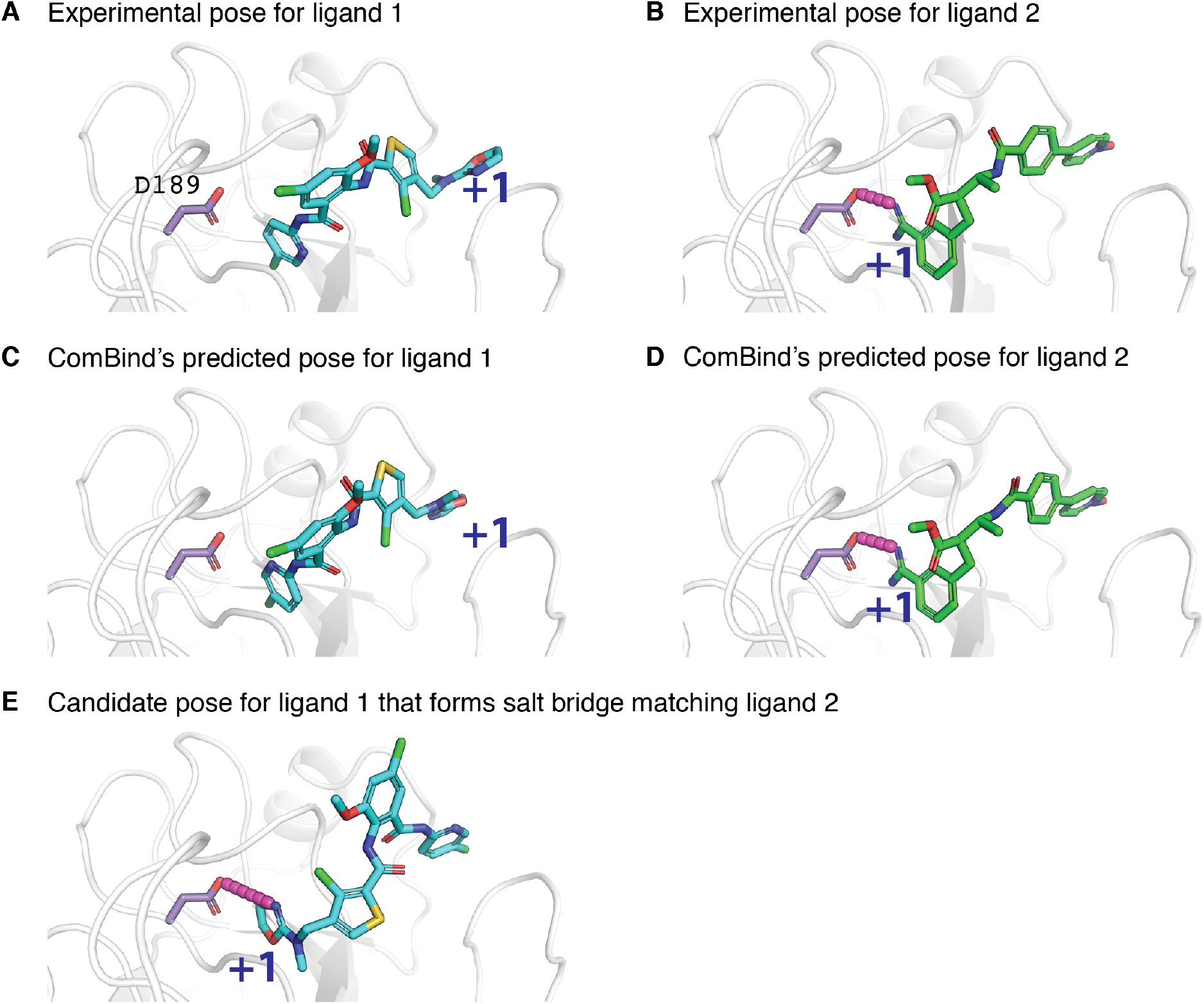
Example of a case where ComBind correctly predicts that ligands form distinct interactions with the protein. We ran ComBind for 20 ligands that bind F10. While most of the ligands have a positively charged group, only some of them position it to form a salt bridge with D189 (e.g., ligand 1, shown in panel A) while others orient it in the complete opposite direction (e.g., ligand 2, shown in panel B). ComBind correctly predicts both binding poses (C, D). (E) One of the candidate poses for ligand 1 forms the same salt bridge as ligand 2. ComBind correctly avoided choosing this pose, even though choosing it would have led to more similar interactions between ligands.

## Supplementary Tables

**Supplementary Table 1:** Performance of Glide SP and Glide XP on our benchmark set. The data presented in this table does not include ligands that share a substantially sized chemical scaffold with the ligand present in the experimental structure used for docking. Including such ligands increases the success rate for both Glide SP and Glide XP (to 49%, 53%, 47%, respectively).

| # Ligands | Is the top-ranked pose correct? |  |  | Is any candidate pose correct? |  |  |
| --- | --- | --- | --- | --- | --- | --- |
|  | *SP* | *XP* | *IFD* | *SP* | *XP* | *IFD* |
| 327 | 44% | 45% | 40% | 81% | 63% | 81% |

**Supplementary Table 2:** Structural data used for benchmarking ComBind. From left to right, columns represent: Protein family, protein name, Uniprot ID, ChEMBL target ID, number of ligands, number of ligands that do not share a scaffold with the ligand present in the experimental structure used for docking, and number of ligands that do not share a scaffold with the ligand present in the experimental structure used for docking and have at least one correct candidate pose. The right-most column corresponds to the number of ligands included in our benchmarks for each target protein.

| PROTEIN FAMILY | PROTEIN | UNIPROT | CHEMBL | # TOTAL LIGANDS | # DIVERSE LIGANDS | # DIVERSE LIGANDS WITH AT LEAST ONE CORRECT CANDIDATE POSE |
| --- | --- | --- | --- | --- | --- | --- |
| GPCR | 5-HT<sub>2B</sub> | P41595 | CHEMBL1833 | 5 | 5 | 5 |
|  | β<sub>1</sub>AR | P07700 | CHEMBL213 | 11 | 6 | 6 |
|  | β<sub>2</sub>AR | P07550 | CHEMBL210 | 7 | 4 | 4 |
|  | mGluR5 | P41594 | CHEMBL2564 | 4 | 3 | 1 |
|  | Smo | Q99835 | CHEMBL5971 | 4 | 3 | 2 |
| ION CHANNEL | GluN1/2A | Q05586<br>Q12879 | CHEMBL1907604 | 8 | 6 | 4 |
|  | GluR-2 | P19491 | CHEMBL3503 | 15 | 7 | 6 |
|  | GluK1 | P22756 | CHEMBL2919 | 18 | 18 | 15 |
| TRANSPORTER | DAT | Q7K4Y6 | CHEMBL238 | 8 | 8 | 7 |
|  | SERT | P31645 | CHEMBL228 | 4 | 4 | 4 |
|  | GLUT1 | P11166 | CHEMBL2535 | 2 | 1 | 1 |
| NUCLEAR RECEPTOR | ER | P03372 | CHEMBL206 | 20 | 14 | 12 |
|  | GR | P04150 | CHEMBL2034 | 16 | 10 | 8 |
|  | MR | P08235 | CHEMBL1994 | 12 | 10 | 9 |
|  | AR | P10275 | CHEMBL1871 | 19 | 12 | 10 |
|  | VDR | P11473 | CHEMBL1977 | 20 | 3 | 3 |
| PROTEASE | F2 | P00734 | CHEMBL204 | 20 | 19 | 15 |
|  | F10 | P00742 | CHEMBL244 | 20 | 19 | 12 |
|  | PLAU | P00749 | CHEMBL3286 | 20 | 20 | 19 |
|  | P00760 | P00760 | CHEMBL3769 | 20 | 19 | 16 |
|  | BACE1 | P56817 | CHEMBL4822 | 20 | 19 | 7 |
| **PHOSPHORYLASE** | PYGM | P00489 | CHEMBL4696 | 20 | 5 | 4 |
| **PHOSPHATASE** | PTPN1 | P18031 | CHEMBL335 | 20 | 19 | 8 |
| **TRANSCRIPTION FACTOR** | BRD4 | O60885 | CHEMBL1163125 | 16 | 13 | 7 |
| **CHAPERONE** | HSP90-α | P07900 | CHEMBL3880 | 20 | 16 | 10 |
| **PHOSPHO-DIESTERASE** | PDE10A | Q9Y233 | CHEMBL4409 | 20 | 19 | 17 |
| **RECEPTOR** | $\sigma_1$ | Q99720 | CHEMBL287 | 4 | 4 | 3 |
| **ELATASE** | ELANE | P08246 | CHEMBL248 | 8 | 1 | 1 |
| **REDUCTASE** | DHFR | P00374 | CHEMBL202 | 20 | 20 | 15 |
| **KINASE** | Cdk2 | P24941 | CHEMBL301 | 20 | 20 | 17 |

**Supplementary Table 3:** ComBind is robust to cases where some of the ligands considered have no correct (near-native) candidate pose. Here we show the results of running ComBind for 20 ligands that bind PTPN1. We considered ligands whose binding poses have been determined experimentally, so that we could assess whether the predicted poses are correct. For over half of the ligands, there were no correct candidate poses (likely because these ligands induce a conformational change in the binding pocket). Despite this, ComBind produces more accurate pose predictions than state-of-the art per-ligand docking software. The ligands used in the predictions correspond to those present in the following PDB structures: 1C88, 1C86, 1GFY, 1ECV, 1C83, 1C84, 1L8G, 1KAV, 1BZJ, 1NWL, 1G7F, 1QXK, 1PYN, 1G7G, 1NZ7, 1NNY, 1NO6, 1ONZ, 1NL9, 1ONY.

|  | Ligand |  |  |  |  |  |  |  |  |  |  |  |  |  |  |  |  |  |  |  |
| --- | --- | --- | --- | --- | --- | --- | --- | --- | --- | --- | --- | --- | --- | --- | --- | --- | --- | --- | --- | --- |
|  | a | b | c | d | e | f | g | h | i | j | k | l | m | n | o | p | q | r | s | t |
| Is any candidate pose correct? | Y | Y | Y | Y | Y | Y | Y | Y | Y | N | N | N | N | N | N | N | N | N | N | N |
| Is Glide's predicted pose correct? | Y | Y | Y | Y | Y | Y | N | N | N | N | N | N | N | N | N | N | N | N | N | N |
| Is ComBind's predicted pose correct? | Y | Y | Y | Y | Y | Y | Y | Y | N | N | N | N | N | N | N | N | N | N | N | N |

**Supplementary Table 4:** Ligands used in predictions for the β1 adrenoceptor. From left to right, columns represent: index of ligand (a–k are as shown in **Fig. 3**; xtal denotes the cocrystallized ligand in the protein structure used for docking), name of ligand, mode of action, and PDB ID of the experimental structure.

| Index | Ligand | Mode of action | Structure |
| --- | --- | --- | --- |
| xtal | cyanopindolol | antagonist | 2VT4 |
| a | dobutamine | partial agonist | 2Y00 |
| b | carmoterol | partial agonist | 2Y02 |
| c | isoprenaline | full agonist | 2Y03 |
| d | salbuterol | partial agonist | 2Y04 |
| e | carazolol | inverse agonist | 2YCW |
| f | iodocyanopindolol | antagonist | 2Y CZ |
| g | 4-(piperazin-1-yl)-1H-indole | antagonist | 3ZPQ |
| h | 4-methyl-2-(piperazin-1-yl)quinoline | antagonist | 3ZPR |
| i | bucindolol | antagonist | 4AMI |
| j | carvedilol | inverse agonist | 4AMJ |
| k | methylcyanopindolol | inverse agonist | 5A8E |

**Supplementary Table 5:**
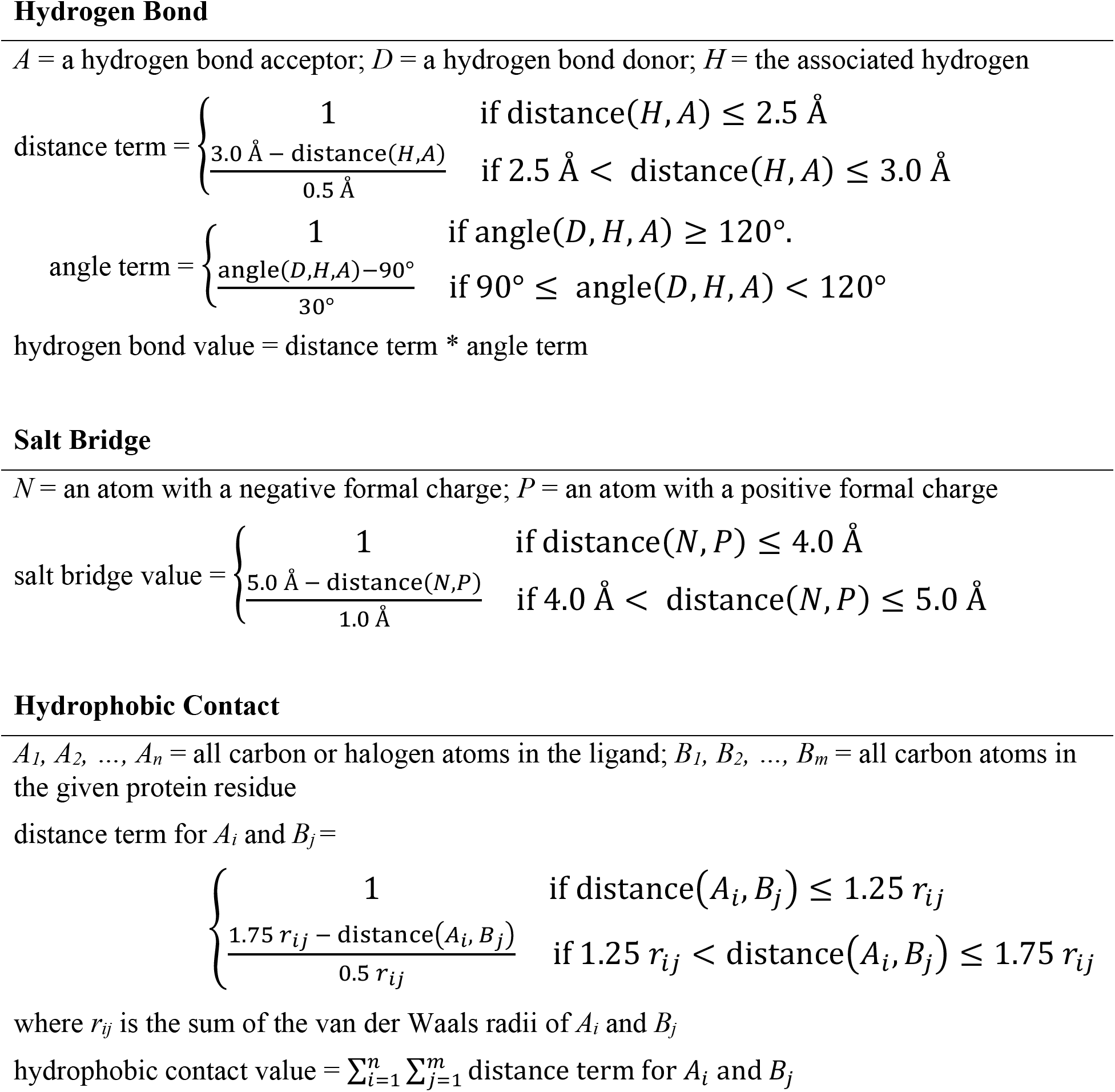
Definitions for the measures used to quantity the presence of each of the three interaction types considered in this study.

**Supplementary Table 6:** Atom types used in maximum common substructure definition. SMARTS pattern and intuitive description of each atom type used when searching for common substructures. Each atom in a molecule is assigned the most specific atom type (lowest in the table) that applies to it.

| SMARTS | Description |
| --- | --- |
| (*) | Any Atom |
| (#1) | Hydrogen |
| (#6) | Carbon |
| (#6; r5; CX4)<br>(#6; r6) | Saturated carbon in 5-member ring<br>Carbon in 6-member ring |
| c1ccccc1 | Carbon-only aromatic ring |
| (CR0) | Carbon not in a ring |
| (#7) | Nitrogen |
| (#7; r5) | Nitrogen in 5-member ring |
| (#8) | Oxygen |
| O=* | Ketone Oxygen |
| (#8; r5) | Oxygen in 5-member ring |
| (#15) | Phosphorus |
| (#16) | Sulphur |
| (#16; r5) | Sulphur in 5-member ring |
| (#9, #17, #35, #53) | Halogens |

